# CapTrap-Seq: A platform-agnostic and quantitative approach for high-fidelity full-length RNA transcript sequencing

**DOI:** 10.1101/2023.06.16.543444

**Authors:** Silvia Carbonell-Sala, Julien Lagarde, Hiromi Nishiyori, Emilio Palumbo, Carme Arnan, Hazuki Takahashi, Piero Carninci, Barbara Uszczynska-Ratajczak, Roderic Guigó

**Author notes:** Corresponding author Correspondence should be addressed to R.G. or B.U.R.

## Abstract

Long-read RNA sequencing is essential to produce accurate and exhaustive annotation of eukaryotic genomes. Despite advancements in throughput and accuracy, achieving reliable end-to-end identification of RNA transcripts remains a challenge for long-read sequencing methods. To address this limitation, we developed CapTrap-seq, a cDNA library preparation method, which combines the Cap-trapping strategy with oligo(dT) priming to detect 5’capped, full-length transcripts, together with the data processing pipeline LyRic. We benchmarked CapTrap-seq and other popular RNA-seq library preparation protocols in a number of human tissues using both ONT and PacBio sequencing. To assess the accuracy of the transcript models produced, we introduced a capping strategy for synthetic RNA spike-in sequences that mimics the natural 5’cap formation in RNA spike-in molecules. We found that the vast majority (up to 90%) of transcript models that LyRic derives from CapTrap-seq reads are full-length. This makes it possible to produce highly accurate annotations with minimal human intervention.

## INTRODUCTION

The processing of eukaryotic RNA molecules is essential for their functionality, with capping and polyadenylation playing key roles. Capping entails the addition of a modified guanine nucleotide (7-methylguanosine) to the 5’ end of the RNA molecule, while polyadenylation involves the addition of multiple adenosine residues to the 3’ end^1^. These modifications provide stability, facilitate export, and ensure proper protein-coding capacity and biochemical activity of non-coding RNAs^2, 3^. Through alternative splice sites, transcription start sites (TSSs), and transcription termination (TTS) or polyA sites, genes generate a diverse range of protein-coding and non-coding RNA molecules^4^. Furthermore, during annotation, the presence of the cap and the poly(A) tail serves as a sequence tag to evaluate transcript completeness^5^.

Understanding the complexity of the transcriptome is crucial for unraveling the principles of gene regulation in contexts like cellular differentiation, organismal development, and disease mechanisms^6–8^. However, current RNA sequencing techniques have limitations that impede the accurate annotation of intact RNA molecules at the transcript level. Long-read RNA sequencing (RNA-seq) methods, like Pacific Biosciences (PacBio) and Oxford Nanopore Technologies (ONT), offer direct analysis of alternative transcript isoforms^9^. Nonetheless, accurately sequencing complete RNA transcripts remains challenging due to drawbacks in library preparation methods, particularly those utilizing SMART (Switching Mechanism At RNA Termini) technology^10–12^. These methods have two notable constraints. Firstly, their reliance on Reverse Transcriptase (RT) template switching can lead to the generation of spurious cDNA products, including false splice junctions and transcript chimera^13–15^. Secondly, none of these methods guarantee the 5’-to-3’ completeness of the sequenced product, resulting in a significant proportion of cDNA 5’ ends falling short of actual TSSs^5, 16^.

Several custom library preparation methods have emerged to address the issue of incomplete transcript termini^17–20^. However, it should be noted that these approaches are often designed for specific platforms and may require additional sample preparation steps, such as efficient ribodepletion. Consequently, their effectiveness in targeting all transcript types may vary^21, 22^. Despite the rapid progress in long-read sequencing methods, the analysis of the resulting data remains challenging. Existing data processing methods, designed primarily for Illumina short-read sequencing, may not be directly applicable to long-read data^23^. Moreover, current strategies for long-read data analysis heavily depend on pre-existing gene annotations, limiting their capability to detect novel, unannotated transcript models^9^. Additionally, available solutions often lack robust quality control strategies for generated annotations, leading to numerous false-positive transcript and gene structures^24^.

Here, we introduce CapTrap-seq, a method that combines the Cap-trapping strategy^25–28^ with oligo(dT) priming to detect 5’capped full-length transcripts. We also present LyRic, a bioinformatics workflow for transcript identification using long-read RNA-seq data. CapTrap-seq and LyRic are being used to produce and process the transcriptome data used in the GENCODE project^29^, and thousands of CapTrap-seq transcript models have already been included in the GENCODE gene set^30^. Additionally, we describe a novel protocol for capping synthetic RNA spike-in sequences, which we use to evaluate CapTrap-seq’s transcript annotation and quantification capabilities.

We have benchmarked CapTrap-seq and other popular library preparation protocols (Teloprime, direct RNA and SMARTer) in a number of human tissues (brain and heart), using both ONT and Pacbio sequencing platforms. This benchmark by itself is a valuable resource by evaluating full-length protocols, showcasing their tested advantages and limitations. We demonstrate that CapTrap-seq compares favorably to other protocols in a number of metrics. In addition, we show that CapTrap-seq produces quantitative estimates of transcript abundances comparable to those obtained using deeper short-read sequence data.

## RESULTS

We first describe the CapTrap-seq protocol and the associated LyRic bioinformatics pipeline. We then benchmark CapTrap-seq against other popular library preparation protocols using the ONT platform in a highly challenging human brain sample. Next, we investigate the impact of the sequencing platform in CapTrap-seq by sequencing the CapTrap brain library, as well as a CapTrap library from a human heart sample, using PacBio Sequel I and Sequel II platforms, in addition to ONT. Finally, we describe a protocol to cap 5’ends of widely used RNA spike-in sequences, and we use them to benchmark LyRic and to evaluate the performance of CapTrap-seq for transcript annotation and quantification.

### CapTrap-seq and LyRic for full-length transcript identification

The CapTrap-seq protocol (Figure 1A) builds upon the previously established Cap-trapping approach^25–28^, but with specific optimizations for long-read RNA sequencing. The protocol begins with the enrichment of polyadenylated transcripts using the anchored oligo(dT) method for cDNA synthesis (Anchored dT and PolyA+ in Figure 1A). After the first-strand synthesis, the initial round of selection for full-length transcripts occurs through the Cap-trapping approach^25^ (Cap-trapping in Figure 1A). Cap-trapping approach was used to address the issue of partial cDNA sequences and to enrich for full-length cDNAs. In this process, the 5’ cap of intact RNA molecules is modified with biotin, enabling the capture of full-length capped RNAs using streptavidin. To isolate cDNA sequences that accurately replicate the 5’ end of the original RNA, an RNase treatment step is employed, cleaving the single-stranded RNA region that connects the cDNA. This step also removes ribosomal RNAs that lack a cap in their native state. The sequential double-stranded linker ligation to single-stranded cDNA (sscDNA) is a highly specific reaction that accurately recognizes cap and poly(A) tail structures while safeguarding cDNA molecules against degradation^26^. The sscDNA strand is released and subjected to the second round of full-length molecule selection through a 5’ and 3’-end dependent linker ligation step, where double-stranded linkers are annealed to the 5’ and 3’ ends of the cDNA molecule (Cap & Poly(A)-dependent Linker Ligation in Figure 1A). The synthesis of the second strand commences with the ligation of universal adapters to the cDNA molecule flanked by both 5’ and 3’ linkers. By employing universal primers, a Long and Accurate PCR (LA-PCR)^31^ method amplifies longer cDNA templates with exceptional fidelity. This approach effectively enriches the presence of full-length cDNA molecules in the resulting libraries to the desired extent (Full-length cDNA library synthesis in Figure 1A). In summary, the CapTrap-seq protocol utilizes two consecutive rounds of full-length transcript selection, focusing on the 5’ cap and poly(A) ends, to accurately identify complete cDNA molecules.

**Figure 1.**
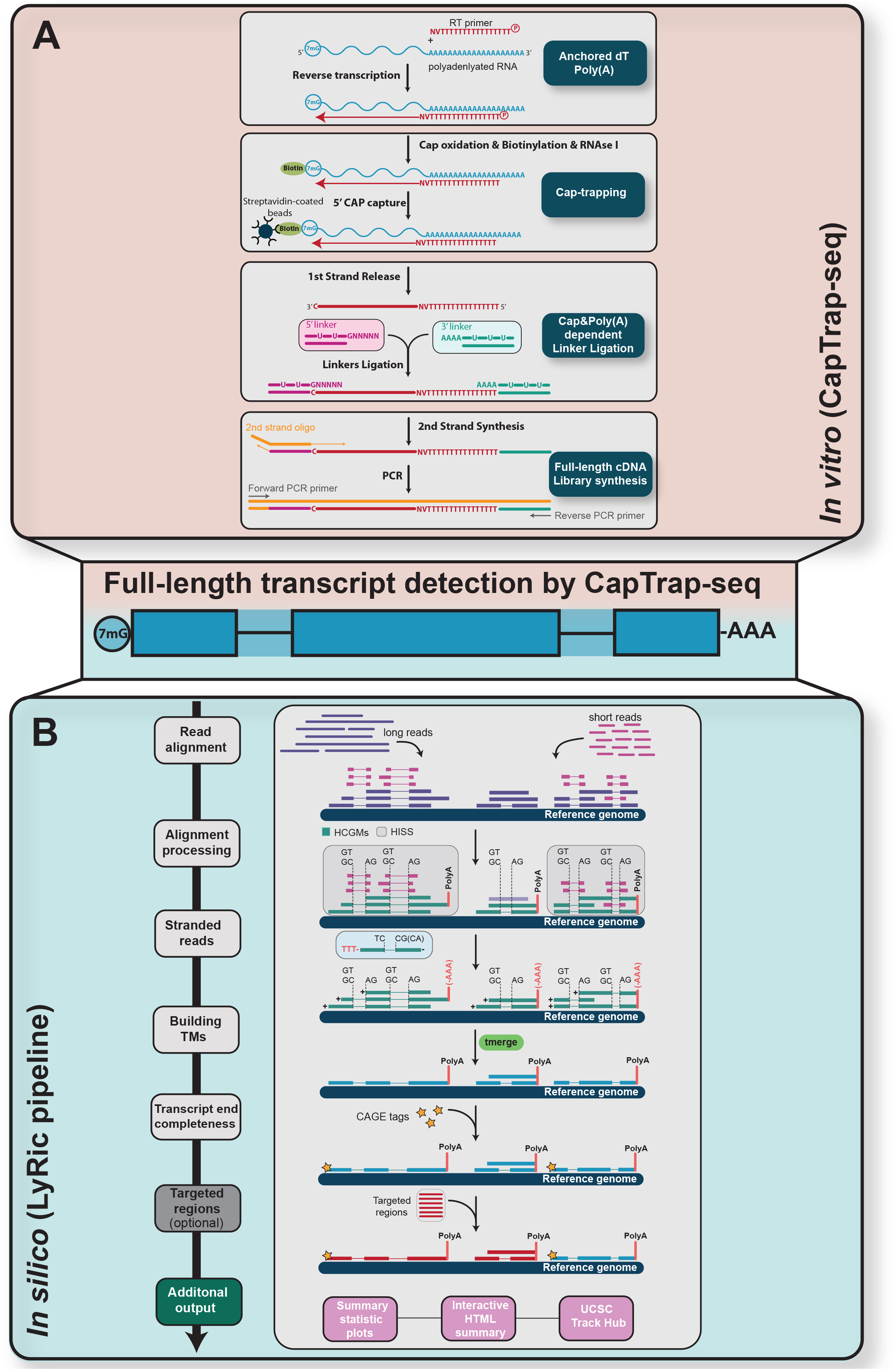
Full-length transcript detection pipeline using CapTrap-seq and Lyric. **(A)** CapTrap-seq experimental workflow. Gray boxes highlight the four main steps of full-length (FL) cDNA library construction: Anchored dT Poly(A)+, CAP-trapping^25–28^, CAP and Poly(A) dependent Linker Ligation, and FL-cDNA library enrichment as described in the text. **(B)** The framework of the LyRic pipeline. The standard long-read data analysis process using LyRic includes five main steps. The read alignment (1) step LyRic maps the long and short (if available) RNA-seq reads to the reference genome using Minimap2^33^ and STAR^48^, respectively. Next, in the alignment processing step (2) the High-Confidence Genome Mappings (HCGMs) and HiSeq Supported reads are identified to be stranded based on their splice sites and poly(A) tail orientation (3). Stranded alignments with compatible intron chains are merged by *tmerge* (see Figure S1 for further details) into non-redundant transcript models in the build TMs step (4). Finally, LyRic evaluates the transcript end completeness using the clipped polyA tail at the 3’-end of the transcript and the support of external CAGE data^34^ (provided by the user) to assess the 5’-end completeness (5). LyRic offers some optional steps to customize the data analysis workflow. By providing a set of non-overlapping capture-targeted regions for each sample in a standard GTF format, LyRic groups features into target types, performs the analysis and generates summary statistics for targeted regions. LyRic offers some additional output including diagnostic plots and UCSC Track Hub. See Materials and Methods for further details.

To streamline the analysis of raw reads obtained from CapTrap-seq experiments and enhance the discovery of novel transcripts while reducing dependence on the pre-existing reference annotations, we have developed LyRic. LyRic is a Snakemake^32^ based bioinformatics pipeline that automates the identification of full-length transcripts from long-read RNA-seq data (Figure 1B) and incorporates quality control at every step. First, LyRic maps long-reads to the reference genome using Minimap2^33^. In the subsequent steps, LyRic performs a filtering process to remove poor-quality alignments. It achieves this by identifying the High-Confidence Genome Mappings (HCGMs) which consist of only canonical and high-quality sequence splice junctions for spliced reads. This filtering step helps eliminate spurious introns that may arise from RT template switching. For unspliced reads, HCGMs require the presence of a detectable, clipped polyA tail (see Materials and Methods). If short-read RNAseq data is available, LyRic performs an additional filtering step on the spliced HCGMs to generate high-quality Hi-Seq-Supported read mappings (HiSS). Importantly, LyRic includes unspliced HCGMs with a poly(A) tail in the transcript model-building process, even though it does not calculate short-read support for them. When short-read data is not utilized to support spliced HCGMs, HiSS reads are equivalent to HCGMs in terms of representation.

The identified HCGMs (or HiSS, if generated) serve as the basis for constructing transcript models using the *tmerge* algorithm (see Materials and Methods). With *tmerge*, compatible reads and transcripts are merged to create non-redundant transcript models. Notably, spliced and unspliced reads are treated separately and never merged together (Figure S1A). To ensure the reliability of the resulting transcript models, *tmerge* incorporates parameters that prevent artificial elongation of intron chains (Figure S1B). Additionally, LyRic addresses the issue of mismapped splice junctions by correcting exon/intron overhangs (Figure S1C). LyRic evaluates the evidence for predicted transcript ends using distinct approaches. The completeness of the 3’ end is assessed based on the presence of an unmapped polyA tail at the transcript’s 3’ end^5^, while the completeness of the 5’ end is evaluated by considering the support from Cap analysis of gene expression (CAGE)^34^ data, if available. CAGE data provides a snapshot of the complete 5’ends of RNA molecules, providing valuable information on transcription start sites. Finally, LyRic offers the capability to compare and potentially merge the newly obtained transcript models with an external gene annotation. This process enables the redefinition of gene boundaries in the new gene set based on a set of overlapping transcripts using the *buildLoci* utility (Figure S1D, see Materials and Methods).

LyRic is a versatile tool for identifying novel transcripts and completing gene structures by detecting missing alternative isoforms. It offers additional settings for analyzing targeted long-read experiments and generates summary statistic plots viewable in an interactive HTML table. LyRic can also create a Track Hub infrastructure to display transcript models in the UCSC genome browser. Although initially developed for CapTrap-seq, LyRic can be used for any unbiased or targeted long-read RNA-seq data. With its modular architecture, LyRic allows for customization and ensures a flexible and tailored data analysis process.

### Benchmarking long-read library preparation protocols

To evaluate the capability of CapTrap-seq in detecting complete and intact transcripts, we conducted a comparative analysis with three popular state-of-the-art library preparation methods (Figure 2A). These methods included (i) SMARTer cDNA synthesis from Takara Bio^10^, (ii) the Oxford Nanopore kit for direct RNA sequencing, and (iii) the TeloPrime approach from Lexogen for full-length cDNA amplification^35^ (Figure S2A). Our study focused on RNA samples derived from the human brain, a tissue known for its high level of transcriptional complexity. For sequencing, we employed the Oxford Nanopore Technologies (ONT) platform, which allows for both cDNA and direct RNA sequencing. To further support the initial predictions made by the High-Confidence Genome Mappings (HCGMs), we also generated short-read RNA-seq data (25 million reads of 125 bp long paired-end reads) using the SMARTer protocol on matched brain samples.

**Figure 2.**
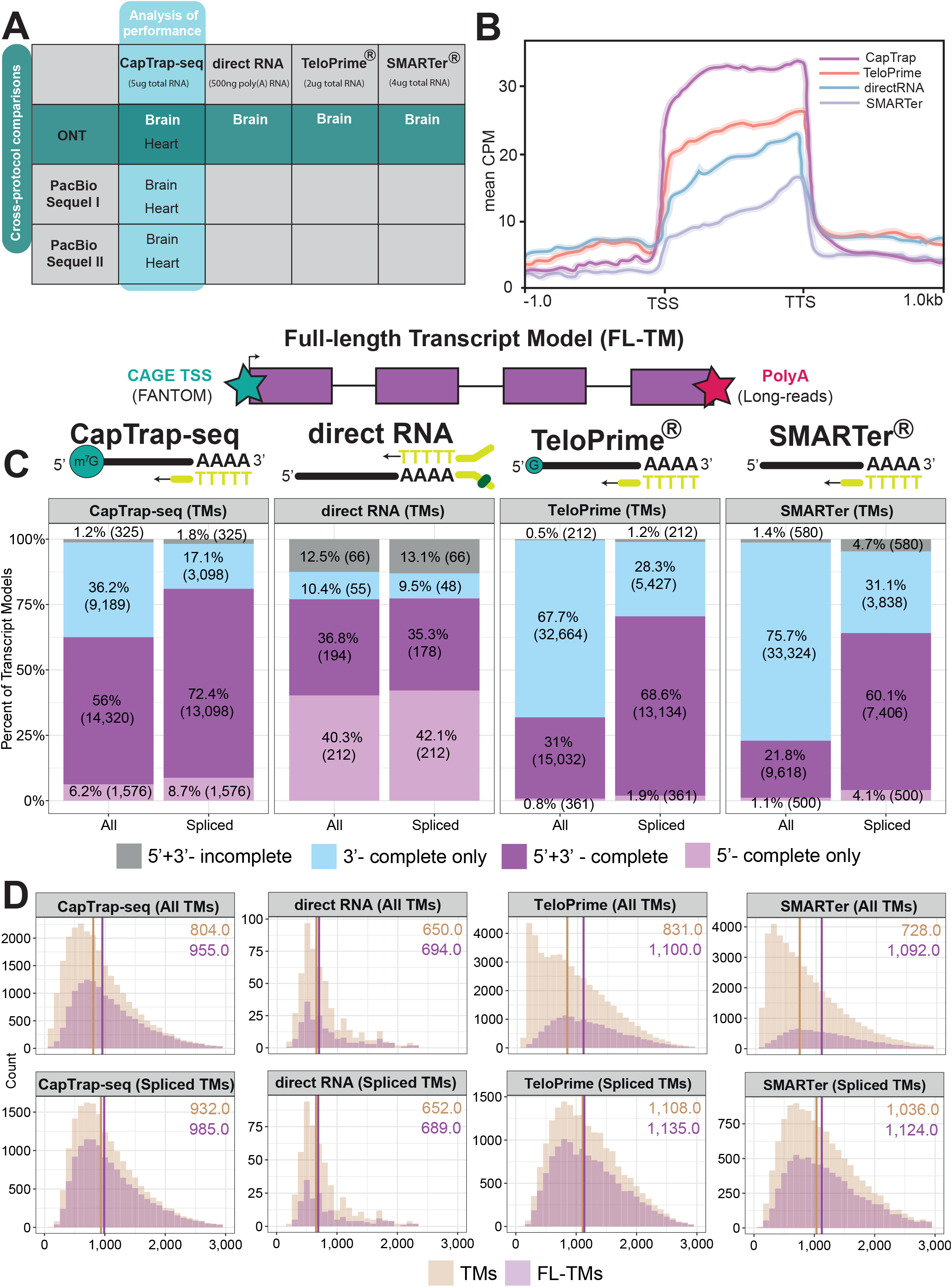
Full-length transcript annotation. **(A)** Two adult human complex transcriptomic samples, brain and heart, were used to perform the cross-protocol and cross-platform comparisons to assess the quality of CapTrap-seq. The horizontal green line indicates the cross-protocol comparisons, including four different sequencing library preparation methods: CapTrap-seq, directRNA®, TeloPrime® and SMARTer®. Whereas, the vertical blue line shows cross-platform comparison using CapTrap-seq in combination with three long-read sequencing platforms: ONT, PacBio Sequel I and Sequel II. **(B)** Read aggregate deptools^49^ profiles along the body of annotated GENCODE genes. **(C)** Detection of full-length transcript models (FL-TMs) among all and spliced TMs with 5′ and 3′ termini inferred from robust (FANTOM5 phase 1/2 robust (*n* = 201,802)) CAGE clusters^39^ and poly(A) tails; Colors highlights four different categories of transcript model (TM) completeness: Grey: incomplete TMs; Sky blue: 3’ complete TMs; Light pink: 5’ complete TMs; Purple: full-length -TMs. **(D)** Length distribution for complete (pink) and incomplete (beige) all and spliced transcript models. The median read length is shown in the top right corner with the color code corresponding to the one specified in the legend.

All protocols, except direct RNA sequencing, produced comparable total numbers of long-reads, with high mapping rates, in particular for CapTrap-seq and TeloPrime (>99%). TeloPrime reads were slightly longer than those produced by CapTrap-seq, which were, in turn, longer than SMARTer, and direct RNA reads (Figure S2B). CapTrap-seq produced the lowest proportion of sequencing errors compared to the other library preparation methods, while this proportion for direct RNA sequencing was the highest (Figure S2C). The lower performance of direct RNA sequencing suggests that this protocol may be particularly sensitive to the integrity of RNA, which tends to be lower when extracted from the brain (RIN=6.5 in the case or our sample). To assess this hypothesis, we sequenced RNA extracted from the human heart (RIN=9.7). Indeed, the number and the length of reads were substantially larger and error rates were lower for heart RNA compared to brain (Figures S2D-E).

We observed differences among the GENCODE biotypes detected by the different technologies when reads were mapped to annotated genes (GENCODE v24). The majority of reads for all protocols overlap protein-coding genes, as expected in poly(A) tail-dependent approaches (Figure S3A). They also represented a similar proportion of long noncoding RNAs (lncRNAs). However, SMARTer produced a comparatively larger fraction of non-exonic reads, including reads mapping to either intergenic or intronic regions. SMARTer and TeloPrime had a larger fraction of reads mapping to poorly understood miscellaneous RNAs, while CapTrap-seq detected more pseudogenes. Notably, CapTrap-seq almost completely eliminated rRNAs that are not capped in native conditions (Figure S3A), thereby minimizing the requirement for an additional ribodepletion step. In terms of nucleotide coverage, approximately 90% of CapTrap-seq’s coverage was in genic regions (exons and UTRs), compared to around 60% for TeloPrime and SMARTer (Figure S3B). Additionally, CapTrap-seq produced the lower percentage of intronic reads (7.3% compared to 23.6% for SMARTer and 17.3% for TeloPrime).

There were also differences in the transcriptional diversity among the protocols, with CapTrap-seq exhibiting a smoother distribution of reads across annotated genes. In CapTrap-seq, approximately 10% of the mapped reads aligned to the top ten genes, whereas this proportion was higher (20-25%) for SMARTer and TeloPrime (Figure S3C). The transcriptional diversity captured by CapTrap-seq closely resembled that of the Illumina short-read dataset, suggesting that it produces a less biased representation of the transcriptome compared to other protocols. Furthermore, CapTrap-seq demonstrated the most uniform read coverage along the length of the GENCODE annotated transcripts (Figure 2B).

In contrast to CapTrap-seq, TeloPrime was designed to detect G at the 5’end of the transcript, not the cap structure itself (Figure S2A). Thus, it will define any transcript starting with G as 5’complete, resulting in potentially numerous false positive predictions. To assess the specificity of CapTrap-seq and TeloPrime, we examined their detection of uncapped synthetic spike-in RNA sequences. We employed two commonly used synthetic RNA spike-in controls: the External RNA Controls Consortium (ERCC) spike-ins with pre-formulated blends of 92 unspliced transcripts to mimic the natural dynamic range of RNA expression and the Spike-In RNA Variants (SIRVs) designed to capture the transcriptomic complexity with 69 different overlapping isoforms grouped in 7 gene modules^36–38^. Despite the lack of a cap structure at their 5’ ends, up to 4% of TeloPrime reads mapped to the synthetic spike-ins, whereas CapTrap-seq, as expected, did not detect any uncapped RNA spike-ins (Figure S4A).

Using LyRic, we processed the raw reads and inferred transcript models (TMs) from them. The initial step involved calling High-Confidence Genome Mappings (HCGMs). CapTrap-seq generated the highest number of HCGMs, with the highest proportion of spliced HCGMs (62.8% compared to 31.7% for TeloPrime and only 19.6% for SMARTer, Figure S4B). The large percentage of unspliced SMARTer reads aligns with the observed high number of non-exonic reads (Figure S3A). While spliced reads are typically more reliable than unspliced reads, as the latter can stem from fragmented transcripts or genomic contamination, polyadenylation in unspliced HCGMs suggests lower genomic DNA contamination risk.We employed the short-read data to support spliced HCGMs and generate HiSS. The support rate was slightly higher for CapTrap-seq (94%) compared to TeloPrime (91%) and SMARTer (88%) when considering only spliced HCGMs. However, when considering the total number of reads in the sample, this proportion was over three times lower for TeloPrime (29%) and SMARTer (17.4%) (CapTrap-seq - 59%, Figure S4B). After running LyRic, SMARTer and TeloPrime produced a greater number of total TMs compared to CapTrap-seq. However, when excluding unspliced models, TeloPrime and CapTrap-seq generated a similar number of spliced TMs, significantly more than SMARTer (Figure S4C).

CapTrap-seq, designed to detect full-length transcripts, outperformed the other methods in identifying complete transcript structures. Using LyRic, which leverages unmapped poly(A) tails at the 3’ end and the proximity to CAGE tags^34^ (FANTOM5 phase 1/2 robust CAGE clusters^39^, N=201,802) for 5’ cap structures to assess transcript completeness, CapTrap-seq identified the highest proportion of 5’+3’ complete transcripts (56% of TMs) compared to Teloprime (31%) and SMARTer (22%) (Figure 2C). This proportion was even higher for spliced transcripts. SMARTer and TeloPrime produced slightly longer full-length transcripts (Figure 2D). CapTrap-seq, on the other hand, showed the largest overlap between CAGE and ENCODE Dnase I hypersensitive sites^40^ (DHS) support, validating the accuracy of detected Transcription Start Sites (Figure 3A).

**Figure 3.**
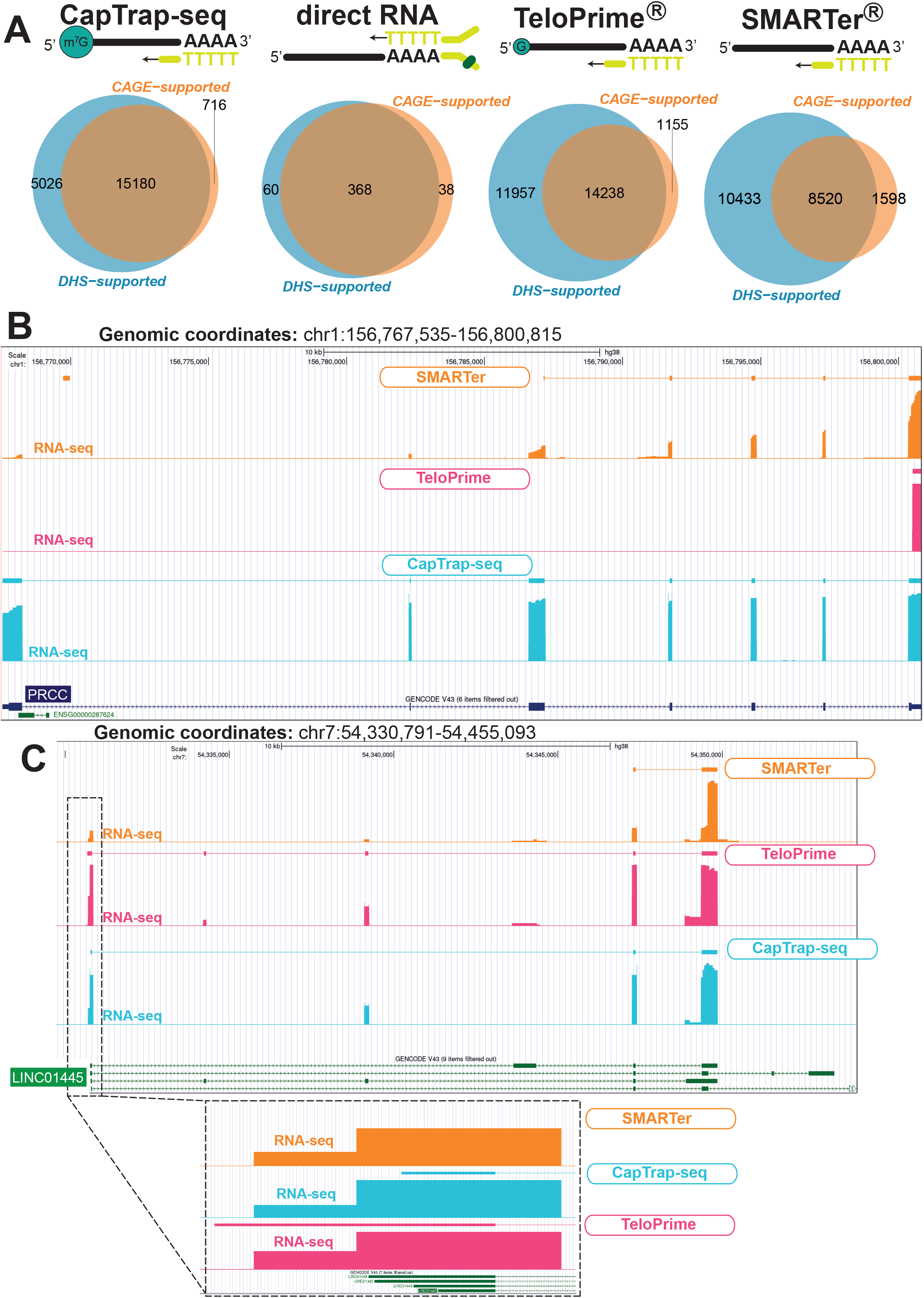
Accuracy of annotated Transcript Models. **(A)** A comparison of TMs supported by FANTOM CAGE tags^39^ (orange) and DNase I hypersensitive sites (DHSs) signals (blue) from the ENCODE data^40^; **(B-C)** Transcript models identified for the *PRCC* protein coding gene (B) and for the *LINC01445* lncRNA gene (C). Colors denote the library preparation method: orange for SMARTer, pink for TeloPrime and blue for CapTrap. The GENCODE models (v43) are shown in navy and green for protein-coding and lncRNA genes/transcripts, respectively. The bigwig files derived from the corresponding long-read RNA-seq data are shown below each transcript model. All TMs are displayed in the dense, while the bigwig uses signal track mode in the full mode in the UCSC genome browser. The dashed line indicates the region that has been enlarged to better visualize the 5’ end of the transcript models.

We next show a few examples of CapTrap-seq efficiency in identifying full-length transcripts. For instance, the *Proline-Rich Mitotic Checkpoint Control Factor* (*PRCC*) gene (genomic and spliced length: ∼33kb and ∼2.1kb, respectively), known for its involvement in pre-mRNA splicing and fusion events with the *Transcription Enhancer Factor 3* (*TEF3*) gene in certain carcinomas, is precisely annotated by CapTrap-seq, capturing the exact TSS, TTS, and exon-intron junctions (Figure 3B). SMARTer, with its bias towards 3’ ends, only detected the 3’ terminal exons of *PRCC*, while TeloPrime and direct RNA methods failed to identify the *PRCC* gene. CapTrap-Seq also excelled in accurately determining transcript ends for long noncoding RNA genes (lncRNAs). This ability is particularly relevant, as lncRNAs remain the largest, yet the most enigmatic component of our genome^41, 42^. In this example, both CapTrap-seq and TeloPrime detect *LINC01445* transcripts, but they identify different isoforms, with TeloPrime extending the 5’ end beyond the GENCODE annotated TSS (Figure 3C).

We also investigated the overlap between the transcript sets generated by the different protocols. Transcripts were considered identical if they had the same intron chains, regardless of differences at the 5’ and 3’ ends. We found significant overlap between SMARTer, TeloPrime, and CapTrap-seq, with approximately 30% of CapTrap-seq and TeloPrime intron chains also detected in SMARTer libraries. Similarly, about 47% of SMARTer intron chains were found in both CapTrap-seq and TeloPrime libraries (Figure S5A). The overlap between the technologies was somehow even larger when considering only full-length transcript models (Figure S5A).

Next, we assessed the performance of the investigated protocols against the GENCODE annotation (v24) at different levels. Firstly, we examined their ability to reliably detect GENCODE genes. TeloPrime and SMARTer identified the highest number of annotated GENCODE genes (17,194 and 16,093, respectively), but a majority (68-73%) of these genes were only partially detected (Figure S5B). In contrast, CapTrap-seq recognized a total of 10,333 genes, with 53% of them being detected from end-to-end. When considering absolute numbers, CapTrap-seq and TeloPrime detected a similar number of full-length genes, which also holds true for full-length spliced transcript models (4,371 vs. 4,329). The partial gene detection is primarily driven by unspliced reads, which give rise to numerous unspliced transcript models and introduce bias in gene expression measurements (Figure S3C).

Secondly, we compared structures of spliced TMs against GENCODE reference. The vast majority of them had intron chains either identical to the GENCODE reference transcript (“Equal”), contained in the reference (“Included”) or different, but associated with known GENCODE genes (“Overlaps”). SMARTer and TeloPrime exhibited similar behavior in this regard, while CapTrap-seq overall reported fewer partial transcripts (“Included”), except for unspliced ones. Moreover, the distribution of transcript categories for all genes in the CapTrap-seq sample resembled that of spliced TMs (both Spliced-All and Spliced-FL), indicating that the influence of unspliced reads (and subsequently TMs) on CapTrap-seq’s performance was minimal (Figure S5C).

Finally, we merged the LyRic models with the GENCODE gene catalog and clustered intergenic and intronic TMs into novel loci. SMARTer produced the highest number of novel loci (13,493 in total, Figure S6A). However, most of these loci consisted of unspliced TMs. When considering only spliced models, both CapTrap-seq and TeloPrime generated twice as many novel loci as SMARTer (Figure S6A). Some of these novel loci represented potential genuine genes, such as a brain-specific, three-exon, full-length TM (Figure S6B). This TM was independently detected by both CapTrap-seq and TeloPrime, and its brain-specific expression was confirmed by GTEx RNA-seq data^43^. Notably, CapTrap-seq demonstrated reliable TM detection even in regions with a more complex transcriptional landscape (Figure S6C). In this particular example, CapTrap-seq, TeloPrime, and SMARTer all identified a novel transcript with brain-specific expression according to GTEx RNA-seq data. However, only CapTrap-seq accurately detected the full-length transcripts, while TeloPrime predicted an additional unspliced alternative transcript that did not fully align with the expression profile observed in GTEx brain data.

### CapTrap-seq performance across sequencing platforms and biological samples

To evaluate the performance of CapTrap-seq in other biological samples and sequencing platforms, we applied CapTrap-seq to a human heart sample using ONT, and to both the heart and brain samples using two PacBio platforms: PacBio Sequel I (PacBioSI) and PacBio Sequel II (PacBioSII)^44^ (Figure 2A). The depth of sequencing for the heart sample on the ONT platform was similar to that of the brain sample, while both the brain and heart samples were sequenced to comparable depth on PacBio. However, the number of PacBio reads, particularly PacBioSI, was lower compared to ONT reads (Figure S7A). The mapping rates and sequencing errors of ONT reads were similar in both heart and brain samples (Figures S7B), but the heart reads were on average longer than the brain reads (Figure S7A).

By employing both PacBio platforms in the brain and heart samples, we achieved overall improvements in CapTrap-seq performance. Both PacBio platforms showed mapping rates that were equal to or higher than those achieved with ONT (Figure S7A). Additionally, the PacBio reads had longer average lengths compared to ONT reads, with minimal error rates (Figure S7B). The PacBio platforms also demonstrated enhanced detection of polyadenylated reads (Figure S7C). In general, we did not observe significant differences in CapTrap-seq performance across platforms in the brain and heart samples, except for potential variations resulting from tissue specificity and potentially higher RNA integrity in the heart.

CapTrap-seq reads mapped to similar gene biotypes and GENCODE annotation (v24), displayed comparable gene biotypes and successful elimination of rRNA across platforms and samples tested (Figure S7D). Moreover, CapTrap-seq exhibited consistent 5’ to 3’ uniform read coverage along transcripts across platforms (Figure S8A) and a similar distribution of reads across genes (Figure S8B).

Next, we employed LyRic to infer TMs from reads in all samples. For the PacBioSII platform, raw spliced HCGMs were employed for building TMs, while for ONT and PacBioSI, we used spliced HCGMs that were further supported by Hi-Seq reads (HiSS) (brain: 25 million, heart: 29 million of Illumina 125bp long paired-end reads), along with raw unspliced HCGMs. The majority of PacBio reads consisted of spliced HCGMs, with up to 95% observed for PacBioSII in the heart (Figure S8C). As a result, the PacBio platforms produced a higher proportion of spliced TMs (Figure S9A).

The PacBio platforms resulted in a higher proportion of full-length transcript models, especially for spliced models (Figure 4A). These transcript models were also, on average, longer compared to those generated by ONT (Figure 4B). Additionally, PacBio sequencing improved the overlap between CAGE and DHS signals, enabling a more accurate TSS detection (Figure S8D).

**Figure 4.**
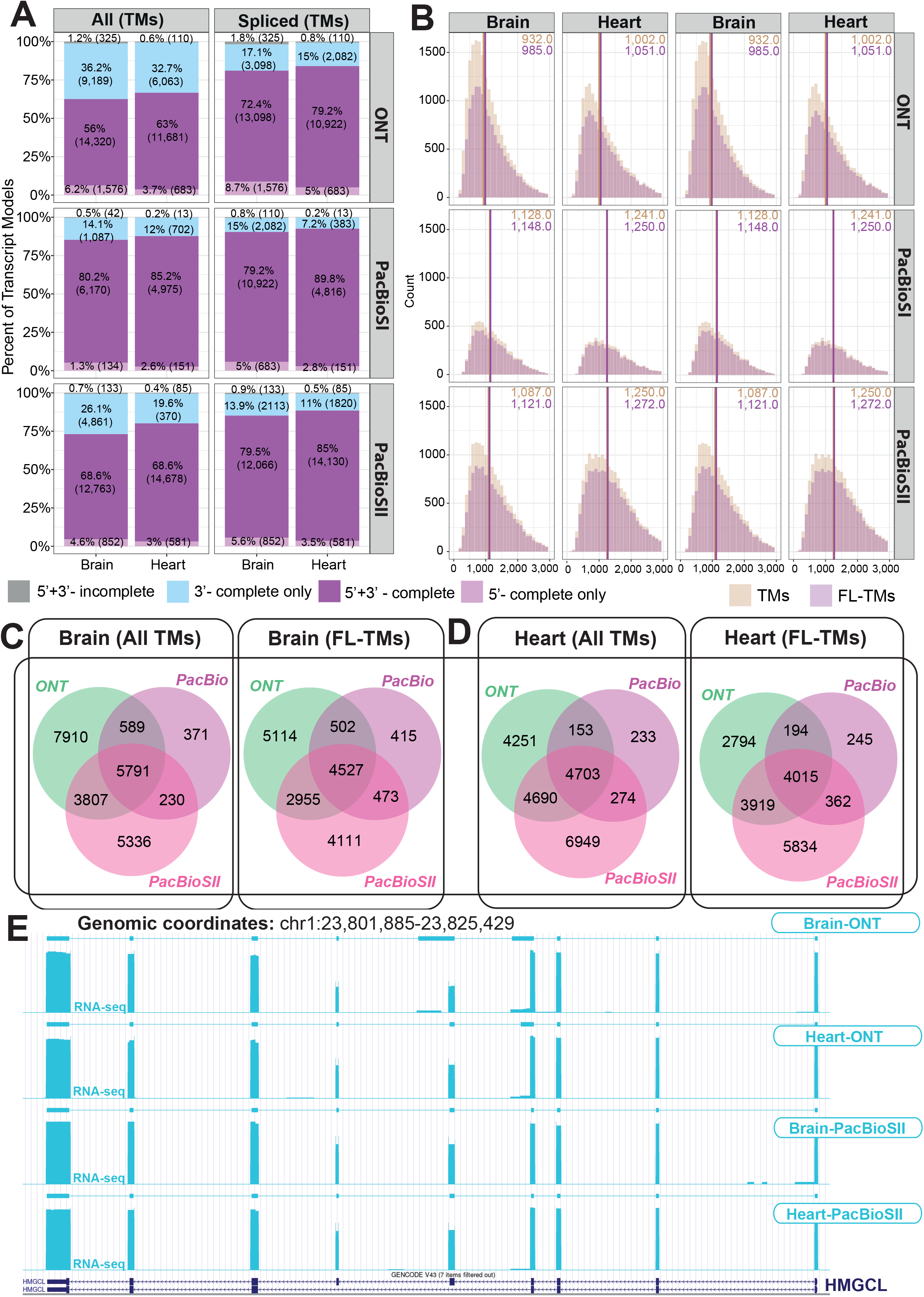
Full-length transcript annotation by CapTrap-seq using different long-read sequencing platforms. **(A)** The proportion of transcript models with different types of termini support as described in Figure 2; **(B)** Length distribution for all (beige) and full-length (pink) transcripts by distinguishing all and spliced TMs; **(C-D)** Detection of all and full-length TMs by ONT and PacBioSI and SII platforms in the brain **(C)** and **(D)** heart samples. **(E)** CapTrap-seq transcript models for the *HMGCL* gene generated using ONT, the PacBioSI and SII platforms. For details see Figure 3.

Therefore, combining CapTrap-seq with PacBio, specifically Sequel II, holds the potential to generate thousands of full-length transcript sequences of manual curation-quality for a wide range of human tissues and conditions.

We observed a significant number of transcript models specific to the ONT and PacBioSII platforms, while the models generated from PacBioSI were mainly a subset of both PacBioSII and ONT models (Figure 4C-D). This suggests that each platform has the ability to detect slightly different RNA molecules, emphasizing the importance of integrating multiple sequencing platforms to achieve a comprehensive profiling of mammalian transcriptomes. Notably, the PacBioSII platform exhibited improved detection of annotated genes compared to PacBioSI and identified a similar number of annotated genes as ONT (Figure S9B).

The performance of CapTrap-seq in combination with PacBioSII is exemplified by the *HMGCL* (*3-Hydroxy-3-Methylglutaryl-CoA Lyase*) gene. We successfully identified this ubiquitously expressed gene in both heart and brain samples using ONT and PacBioSII (Figure 4E). However, PacBioSII enabled more accurate detection of this nine-exon gene (>23kb genomic and ∼1.6kb spliced length). Importantly, the terminal exons, particularly at the 5’ end, were accurately identified by all tested platforms.

Integration of TMs with the GENCODE (v24) annotation using LyRic revealed the presence of numerous novel gene models within intronic and intergenic regions (Figure S9C). Among the sequencing platforms, ONT yielded the highest number of novel genes, including full-length spliced variants. Many of these novel genes remain unannotated even in the latest version of the GENCODE catalog (v43). Notably, an entirely new full-length locus encoding a three-exon transcript, was independently identified by both ONT and PacBio Sequel II (Figure S9D). GTEx^43^ RNA-seq data supported the heart-specific expression of this gene, while it remained undetected in SMARTer and TeloPrime brain-specific experiments. The combination of different sequencing platforms can uncover novel transcript isoforms and further expand the transcriptomic complexity of specific genomic loci. For example, the PacBioSII platform improved the annotation of a previously identified novel locus (Figure S6B) by identifying a previously missing isoform for this gene (Figure S9E).

### Capping RNA spike-in controls for reliable full-length transcript detection

Synthetic controls, such as ERCC and SIRV, are frequently employed to mitigate technical biases in RNA-seq data and analysis. However, these controls lack a cap structure at their 5’ ends, limiting their compatibility with full-length, cap-dependent transcript sequencing methods like CapTrap-seq.

To address this limitation, we have developed a protocol for capping ERCCs and SIRVs (Figure 5A). This protocol mimics the natural 5’ cap formation process by introducing a 7-methylguanosine (m^7^G) cap structure to the synthetic RNA controls through a two-step catalytic process. Firstly, the enzyme guanylyltransferase (GTase) adds GMP derived from GTP to the pp5’N structure located at the 5’ end of the spike-in sequence. Then, the RNA (guanine-N7) methyltransferase (N7MTase) adds a methyl group to the guanine (derived from the added GMP) at the N7 position.

**Figure 5.**
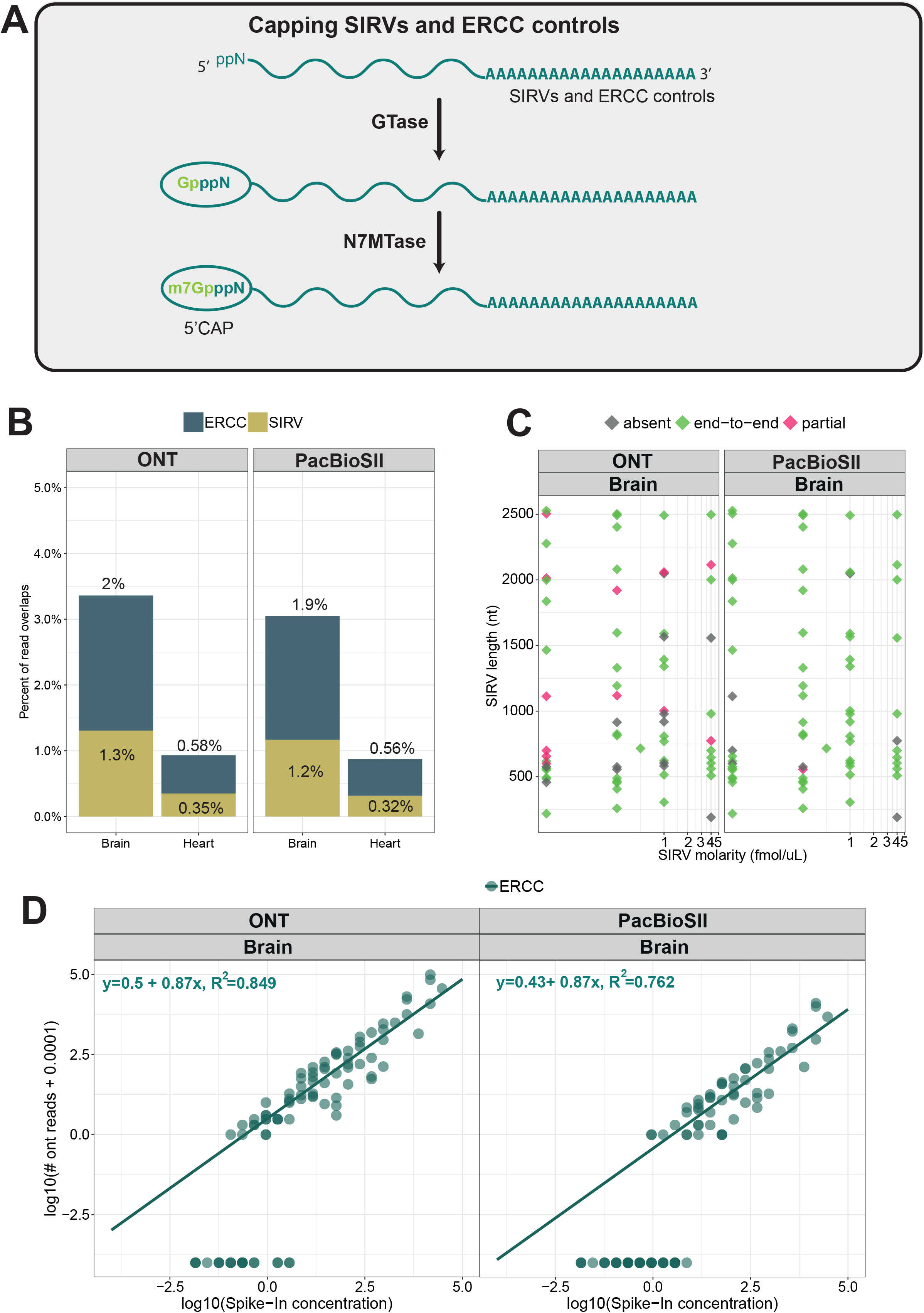
Capping of SIRV and ERCC controls. **(A)** Two-step enzymatic strategy for adding a cap structure at the 5’ends of uncapped RNA spike-in controls; **(B)** Detection rate of SIRV (yellow-green) and ERCC (navy) synthetic controls in the heart and brain samples; **(C)** Detection of SIRVs as a function of initial concentration and length. Three main detection levels have been distinguished: end-to-end (green), partial (red), not detected/absent (gray) **(D)** Correlation between input RNA concentration and raw read counts for ERCC spike-ins in the brain sample. Each point represents a synthetic ERCC control. The green line indicates a linear fit to the corresponding dataset. The parameters are given by the equation on the top.

Both ERCC and SIRV spike-ins, which were modified with newly synthesized 5’ cap structures, were introduced to the total RNA samples prior to the CapTrap-seq library preparation. Subsequently, CapTrap-seq samples from the heart and brain, containing the capped ERCC and SIRV spike-ins, were sequenced using both the ONT and PacBioSII platforms (Figure 10A). The detection of both ERCC and SIRV capped controls was successful in both samples, as illustrated in Figure 5B. It is worth noting that CapTrap-seq did not detect the native, uncapped RNA synthetic sequences (Figure S4A), demonstrating its specificity. However, as depicted in Figure 5B, the detection rate of capped RNA spike-in controls varies depending on the RNA dynamic range of each sample (Figure S8B), with the brain sample demonstrating a higher detection rate compared to the heart sample.

We employed the SIRVs to evaluate the performance of CapTrap-seq in identifying full-length transcripts. CapTrap-seq demonstrated the ability to accurately detect most SIRVs end-to-end, particularly when utilizing the PacBioSII platform (Figure 5C, S10B), regardless of their length and concentration. However, the ONT platform faced more challenges in detecting longer and less abundant transcripts. Next, to assess the quantitativeness of CapTrap-seq, we employed the ERCC spike-ins at known concentrations (Figure 5D, Figure S10C). Our findings revealed that CapTrap-seq was capable of detecting molecules at an approximate concentration of 1.05 x 10^-2^ copies per cell (see Materials and Methods for details). Furthermore, we observed a clear linear relationship between ERCC concentrations and read counts, providing strong evidence that CapTrap-seq can be effectively utilized for transcript quantification. Quantification estimates were more accurate when utilizing the ONT compared to the PacBioSII platform, which could be attributed to the higher throughput of the ONT sequencing (Figure S7A and S9B).

By using SIRVs with well-defined transcript structures, we evaluated the ability of the LyRic pipeline to reconstruct transcript models. LyRic achieved high precision and recall across all investigated structural levels (from single base resolution to the entire transcript structure), especially on the PacBioSII platform. However, ONT models had lower performance, particularly in recovering complete transcript structures (Figure S10D). Incorporating spliced HCGMs supported by short-read RNA-seq data (HiSS) in the LyRic pipeline, improved ONT’s performance (Figure S10E).

## DISCUSSION

In this study, we introduce CapTrap-seq, a library preparation protocol designed to address the issue of incomplete transcript termini in long-read RNA sequencing methods. CapTrap-seq is an open-source, platform-agnostic, and off-the-shelf approach that aims to produce high-confidence full-length transcript models at high-throughput scales. By filtering out uncapped nucleic acids, it effectively mitigates the risk of genomic DNA and rRNA contamination.

Additionally, we present the LyRic pipeline, a fully automated long RNA-seq analysis and gene annotation workflow. LyRic was developed to meet the pressing need for high-quality and high-throughput annotation methods for complex transcriptomes. In contrast to other tools^24^ LyRic does not depend on pre-existing reference annotations, enabling the discovery of novel genes in intergenic regions and the identification of missing alternative isoforms of annotated genes. While we present both of them together, CapTrap-seq and Lyric are not connected, and they can be used independently.

The performance of LyRic and CapTrap-seq, either in combination or as standalone methods, has been evaluated by the Long-read RNA-seq Genome Annotation Assessment Project (LRGASP) Consortium^24^. The LRGASP analysis revealed that CapTrap-seq outperforms other tested protocols in detecting canonical splice junctions and enables reliable transcript reconstruction, as demonstrated by its performance on spliced SIRVs. Additionally, when combined with ONT sequencing, CapTrap-seq exhibits the lowest irreproducibility and highest consistency among other long-read library preparation methods, particularly those using template switching. It ranks second only to the use of RSEM^45^ on short-read Illumina sequencing. In the LRGASP comparison, LyRic is the only pipeline that does not rely on reference annotation guidance. While this leads LyRic to predict a lower number of transcripts compared to other pipelines that rely on annotation, it also enables LyRic to identify a larger number of novel and reliable full-length transcript models. It is important to note that, similar to other evaluated pipelines, LyRic is influenced by platform-specific limitations. In general, Lyric provides more reliable transcript models when used in conjunction with PacBio than with ONT.

In addition to CapTrap-seq, we developed a novel enzymatic capping strategy for synthetic RNA spike-in controls. This strategy enables efficient capping of the 5’ end of both spliced SIRV and unspliced ERCC controls, regardless of their length and initial concentration. By employing these capped spike-in synthetic controls, we were able to evaluate the sensitivity, quantitativeness, and accuracy of the CapTrap-seq method. These capped spike-in controls, which mimic the dynamic range of RNA expression and alternative splicing, have been utilized in the LRGASP project, contributing to the precise annotation of full-length transcript models^24^.

While CapTrap-seq offers several advantages, it is important to consider its limitations. Firstly, a significant amount of starting RNA (5μg) is currently required, but efforts are underway to improve the protocol’s efficiency with smaller amounts of material. Secondly, CapTrap-seq is a relatively complex laboratory procedure, and its multi-stage nature may promote the detection of shorter RNA molecules compared to standard library preparation methods such as SMARTer. However, no length-specific bias was observed for the ERCC and SIRV controls, which are up to 2.5kb long, making CapTrap-seq suitable for analyzing the majority of human RNAs, which have a median length of 2.3kb^46^. Thirdly, CapTrap-seq currently focuses primarily on identifying polyadenylated RNA molecules. Adapting the protocol for the analysis of non-polyadenylated transcriptomic fractions would require substituting the oligo-dT probes and the 3’ linker with custom adapters^17, 20^, although this modification would need further testing.

In conclusion, long-read RNA sequencing, coupled with appropriate data analysis tools, has the potential to produce for the first time highly accurate full length transcript maps of eukaryotic genomes, without the need for human curation. This is particularly important as projects to sequence the genomes of all eukaryotic species on Earth are underway^47^. We believe CapTrap-seq could play an important role in generating the high quality transcript data needed to produce accurate annotations of these genomes.

## SUPPLEMENTARY FIGURES

**Figure S1.**
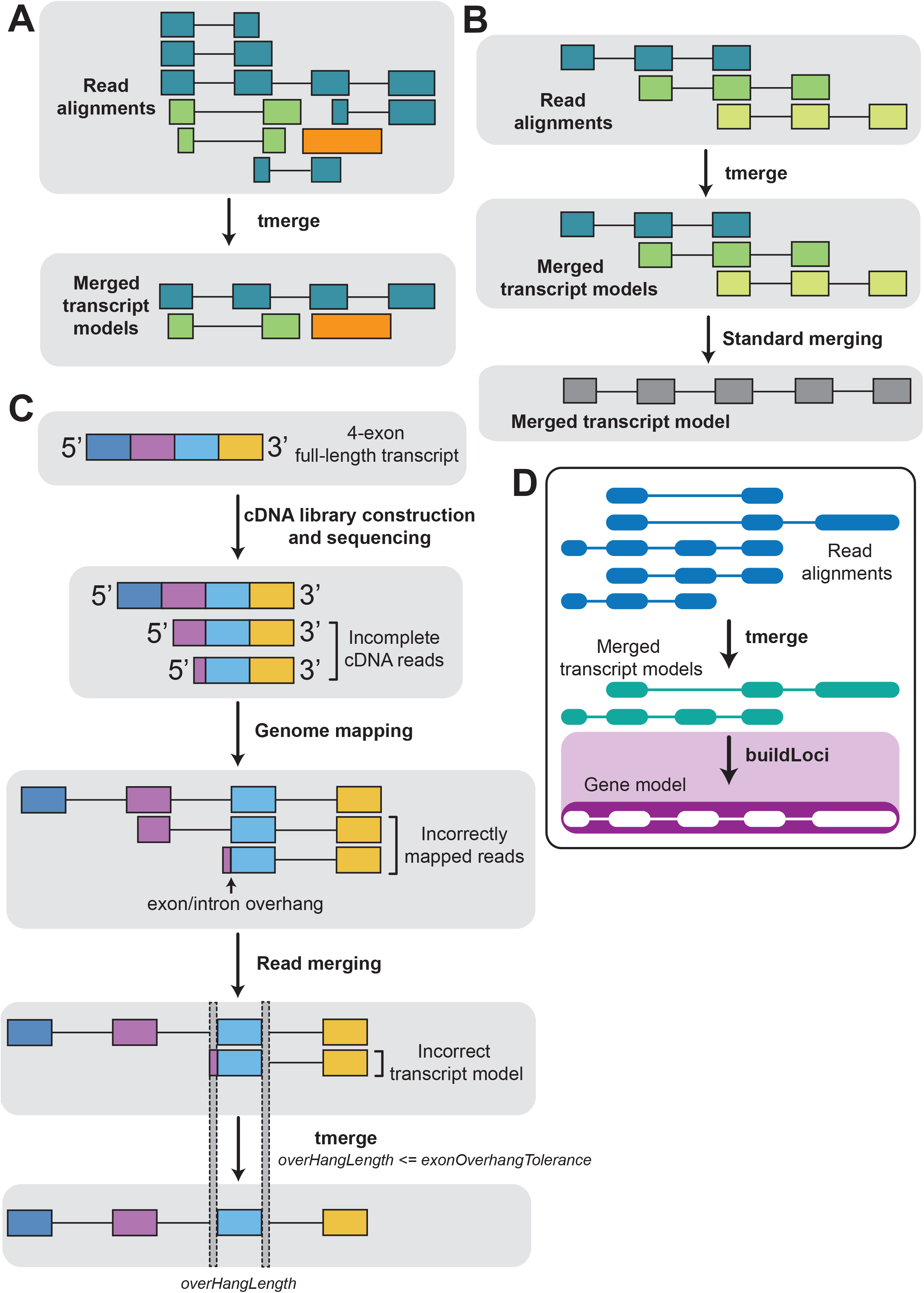
Building accurate TMs from the long-read alignments. **(A)** Transcript model building using *tmerge*. *tmerge* compares read-to-genome alignments (or transcript structures) present in the input and attempts to reduce transcript redundancy by merging compatible input long-reads into non-redundant set of transcript models. The program treats spliced and unspliced reads separately (*i.e.*, those are never merged together). If both structures are spliced, they are deemed compatible if: (1) at least one of their exons overlap on the same genomic strand, (2) either their intron chains are equal, or one is an exact subset of the other, (3) there is no overlap between an exon of one structure and an intron of the other; **(B)** In contrast to standard merging, *tmerge* will never artificially extend intron chains. See merging condition #2 in panel A; **(C)** *tmerge* offers several parameters that allow to fine-tune the TM generation process and increase the quality of output TMs. Setting the exonOverhangTolerance option to a positive integer can correct mismapped splice junctions that sometimes occur when aligning very short, error-rich terminal read exons (for details see https://github.com/guigolab/tmerge). **(D)** *BuildLoci* is a utility to build gene loci (i.e., sets of overlapping transcripts) out of a transcript set produced by *tmerge*. For further details see https://github.com/julienlag/buildLoci.

**Figure S2.**
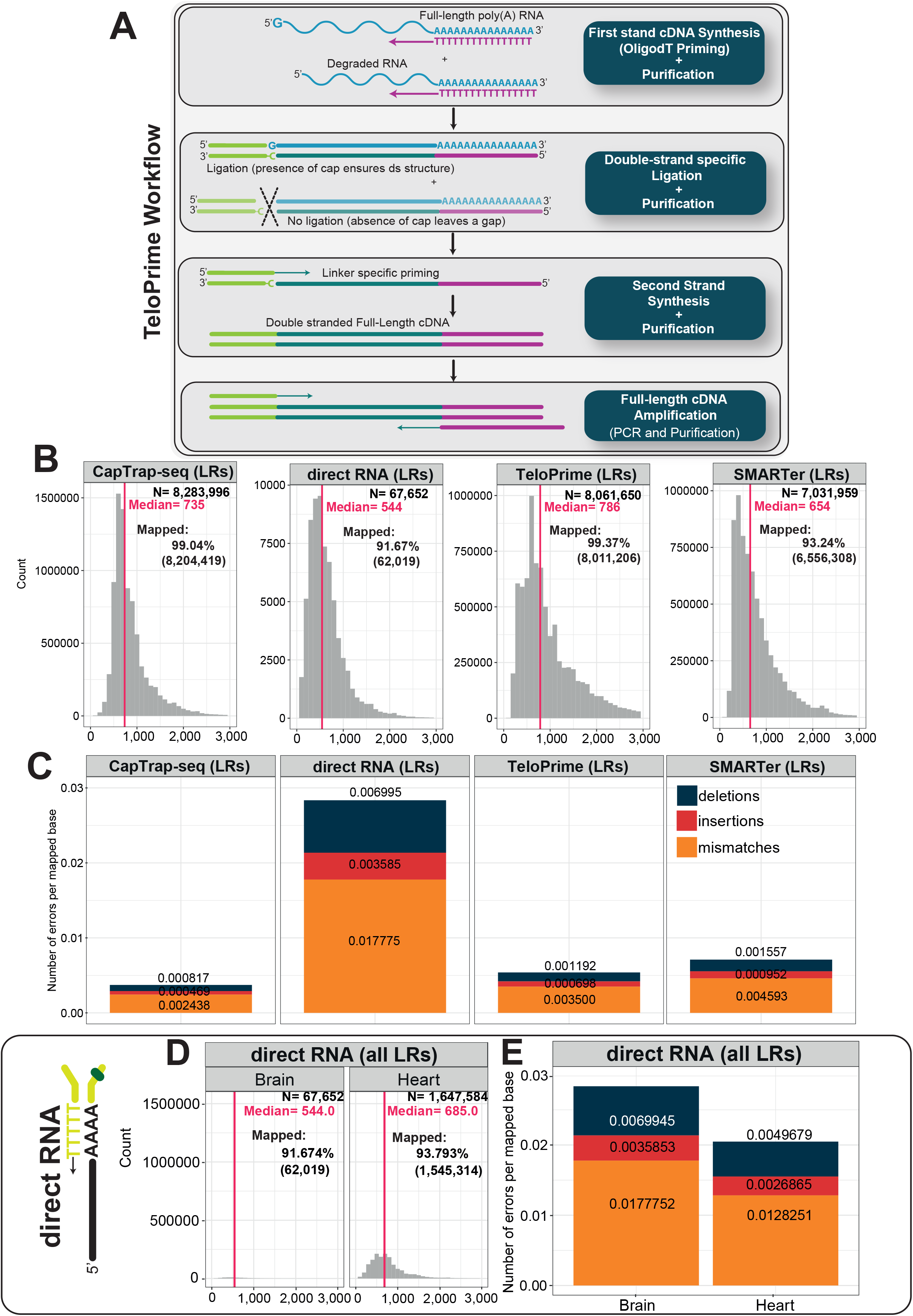
The TeloPrime workflow and the initial read quality check. **(A)** Schematic overview of the TeloPrime Full-Length cDNA Amplification V2 workflow according to materials provided by Lexogen company. **(B)** Length distribution of raw long-read ONT reads for each protocol. The total number of reads (N), median read length (red and vertical line) and the mapping rate are shown in the top right corner. **(C)** The number of errors per mapped bases (error rate) across sequencing library preparation protocols (CapTrap-seq, directRNA®, TeloPrime® and SMARTer®) in the human brain sample. Colors denote the type of error: deletions (black), insertions (red), mismatches (orange); From **D** to **E** metrics for directRNA ONT sequencing in the human brain and heart samples as described above:

**Figure S3.**
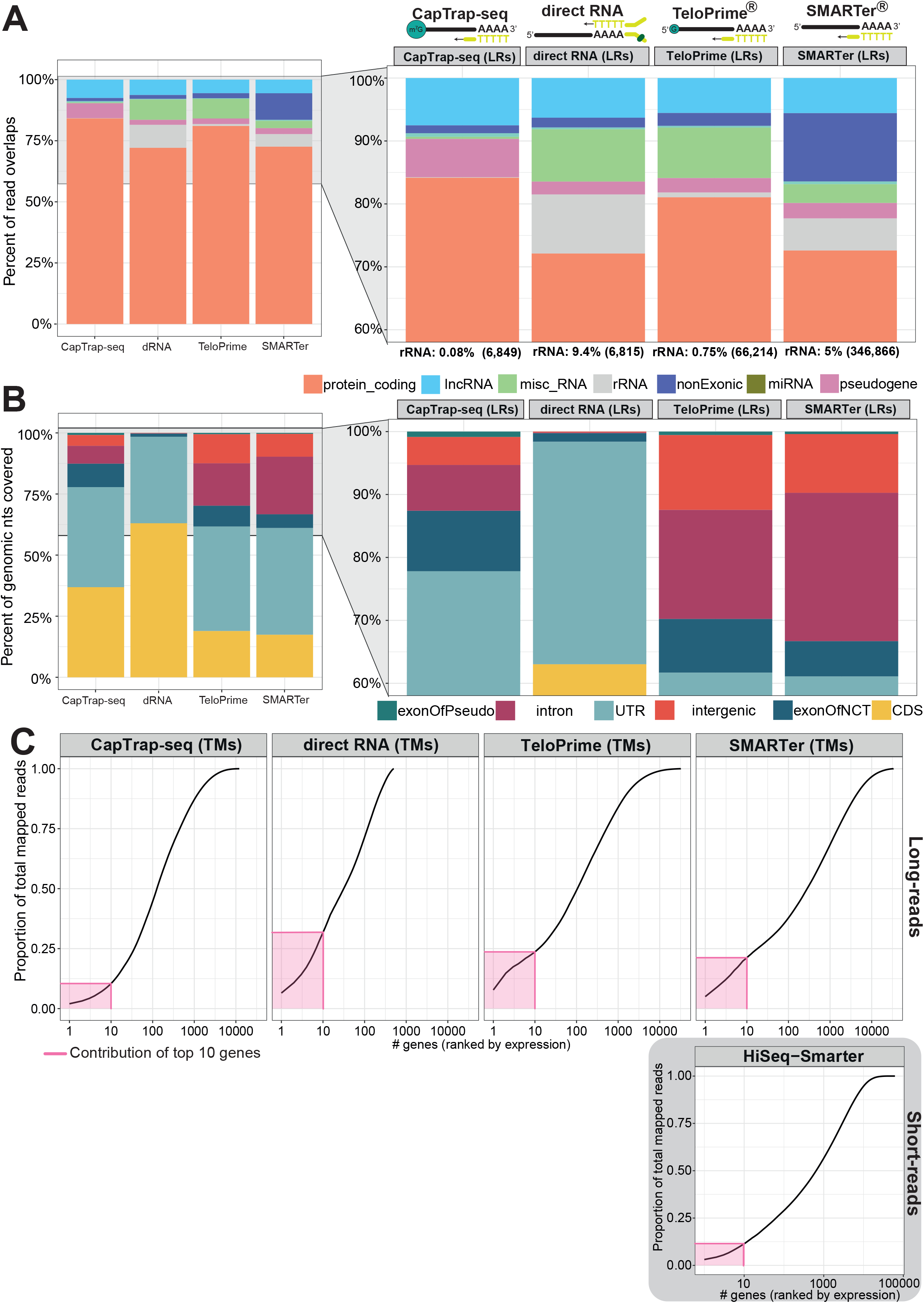
Read quality analysis for the brain sample. **(A)** The GENCODE gene biotype detection by ONT sequencing of four tested libraries. The stacked bar plots report the percentage of raw read overlaps with annotated GENCODE genes (v24) of given biotype. Colors denote different GENCODE gene biotypes: protein-coding genes (orange), ribosomal RNAs (light gray), pseudogenes (pink), miscellaneous RNA (green), miRNAs (olive green), non-exonic (dark blue), lncRNA (sky blue). The proportion of raw reads mapping to rRNA along with absolute raw read numbers in parenthesis is given below each bar plot; **(B)** The proportion of nucleotides covering different genomic partitions. Colors highlight different region types: exon of pseudogene (green), intron (pink), UTR (blue), intergenic (orange), exon of noncoding transcript (navy) and CDS (yellow); **(C)** Cumulative transcript curve showing the proportion of mapped reads (Y axis) captured by the top n more expressed genes (x-axis). The pink area represents the proportion of reads captured by the top ten gene expressors.

**Figure S4.**
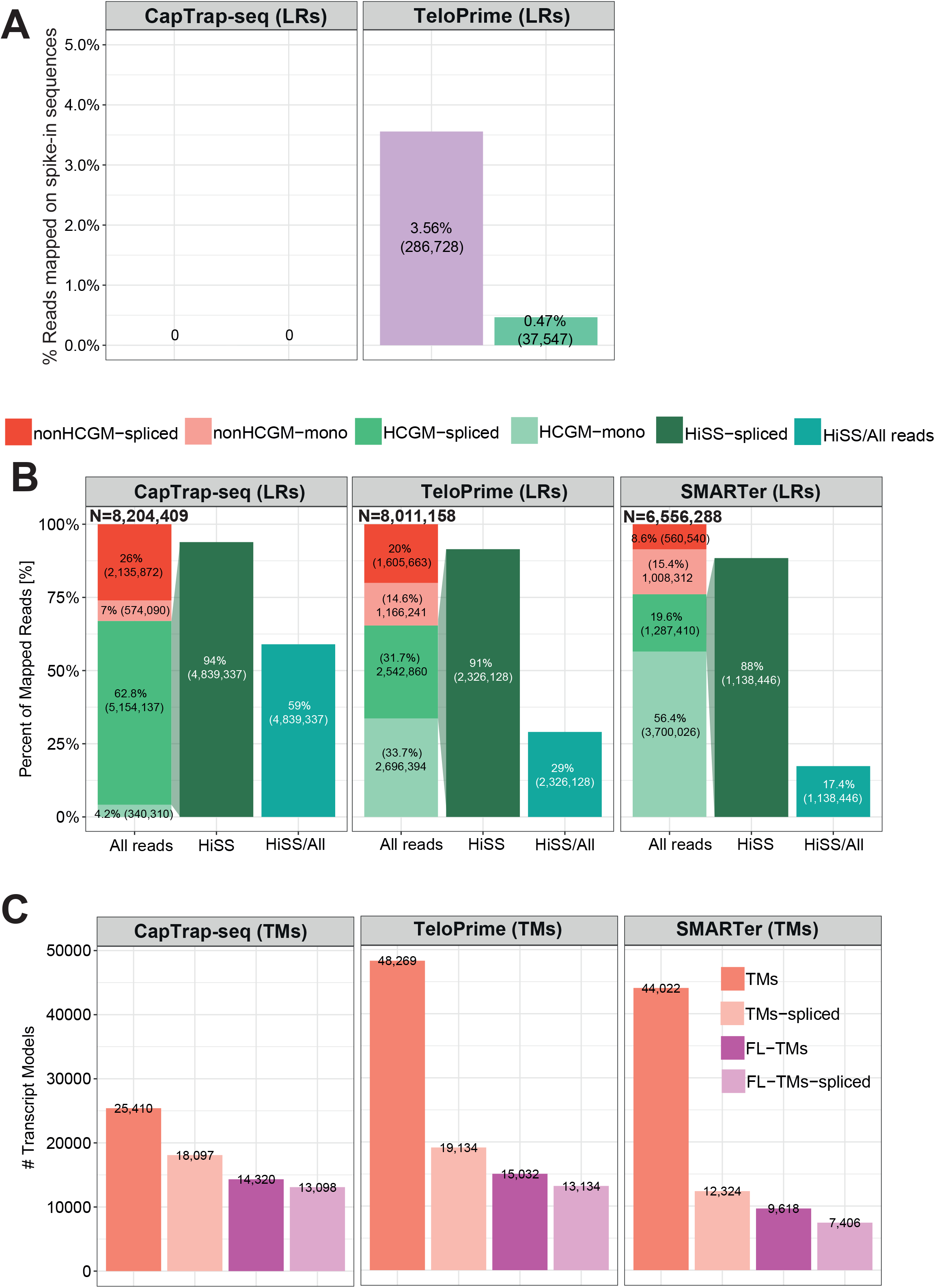
Building transcript models for the brain sample. **(A)** Proportion of ONT reads mapping to the ERCC (lilac) and SIRV (green) synthetic RNA spike-ins in the CapTrap-seq and TeloPrime brain samples. **(B)** The proportion of detected High-Confidence Genome Mappings (HCGMs) and Hi-Seq-Supported read mappings (HiSS). Six main classes include: unspliced HCGMs (light green), spliced HCGMs (dark green), unspliced non-HCGMs (light red), spliced non-HCGMs (dark red), HiSS (dark green) and proportion HiSS to the total number of reads (HiSS/All, turquoise). For a detailed description of how HCGMs and HiSS reads are identified, see the text. The total number of reads (N) after removing ultra-short exons during bam to bed conversion. **(C)** Bar plot showing the number of different TM types, including all TMs (red), spliced TMs (coral), full-length TMs (purple) and full-length spliced TMs (lilac);

**Figure S5.**
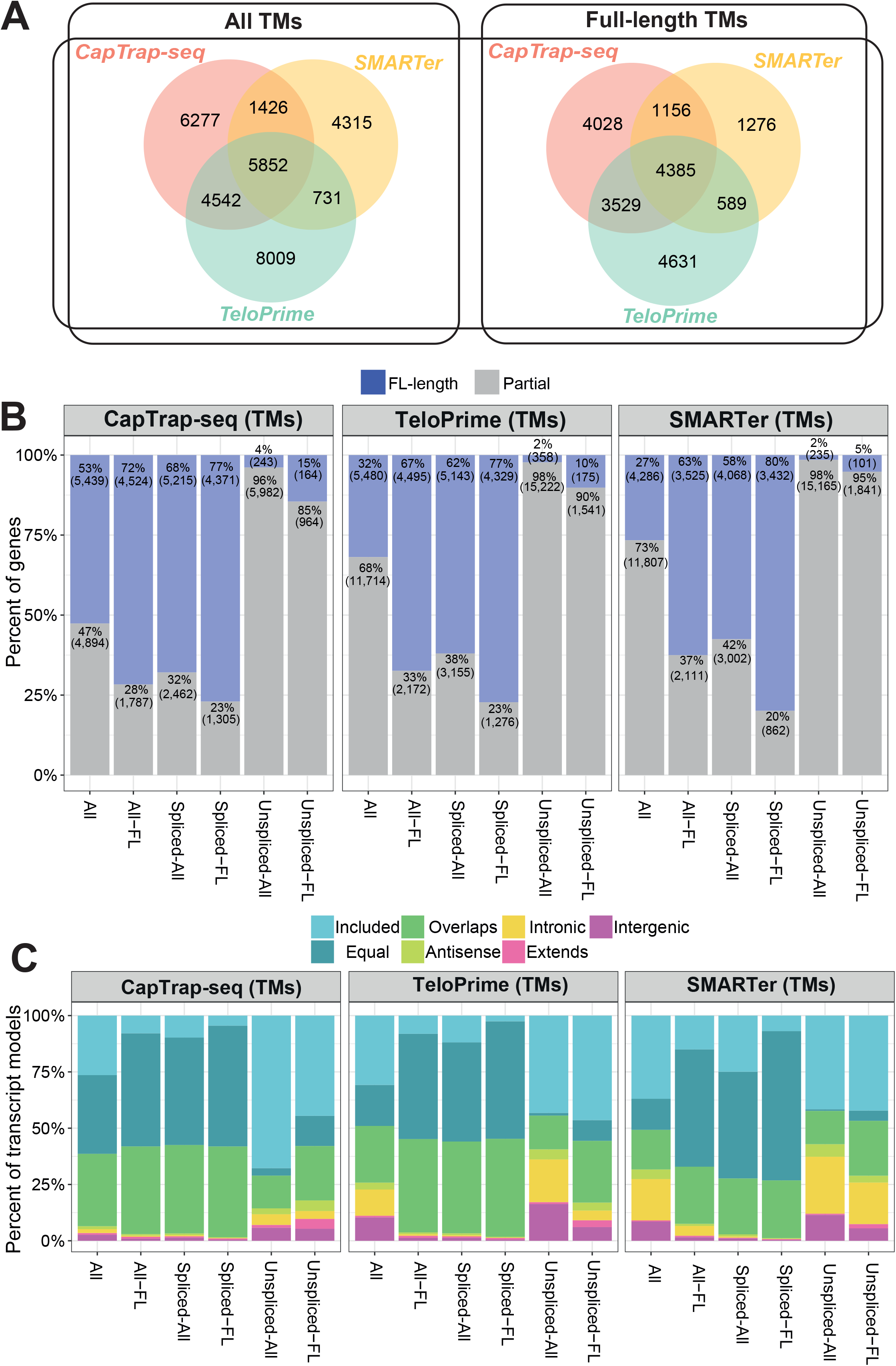
Protocol specificity and gene detection rates. **(A)** 3-way venn diagrams showing the number of identical TMs (all and full-length) detected by three library preparation methods: CapTrap-seq (red), TeloPrime (green), and SMARTer (yellow). We considered two transcripts identical if they shared identical intron chains, irrespective of differences at the 5’ and 3’ termini. **(B)** Proportion of GENCODE (v24) genes fully (end-to-end, blue) or partially (gray) detected by six different classes of identified TMs: all TMs (All), only full-length TMs (All-FL), all spliced TMs (Spliced-All), only spliced full-length TMs (Spliced-FL), all unspliced TMs (Unspliced-All), and only unspliced full-length TMs (Unspliced-FL); **(C)** Comparison of detected TM structures in the human brain samples against GENCODE (v24) annotation. The y-axis represents the number of unique TM counts. The color code describes novelty compared with GENCODE for different TM types. The classes are simplified versions of gffcompare^50^ transcript classification code.

**Figure S6.**
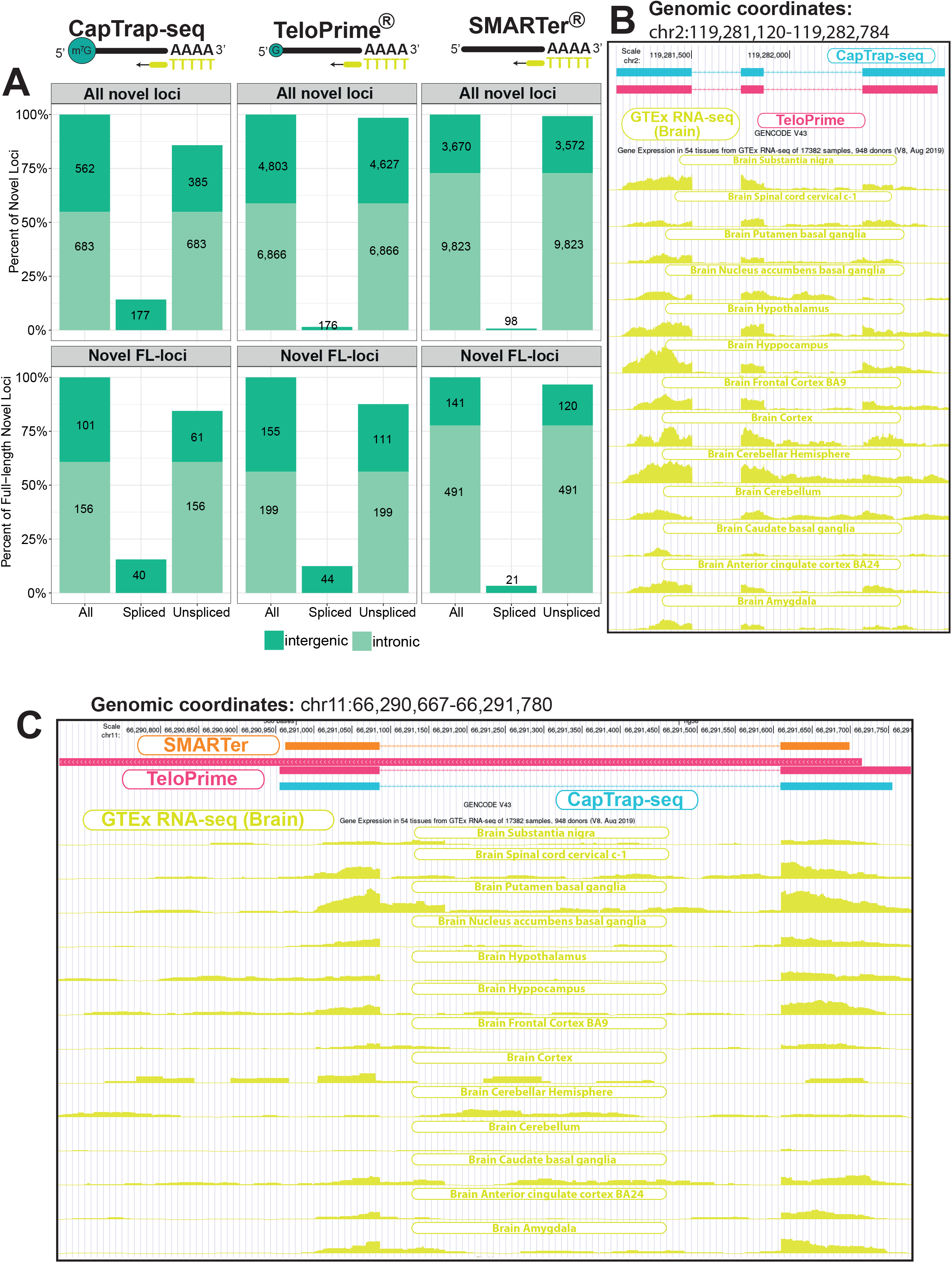
Identification of novel genes by different library preparation protocols. **(A)** The proportion of all novel (top) and novel full-length (bottom) intergenic (dark green) and intronic (light green) loci (all, spliced and unspliced) by tested library preparation protocols. **(B)** and **(C)** Two examples of novel (unannotated in GENCODE v43), full-length spliced loci with brain-specific expression as evidenced by GTEx^43^ RNA-seq data coverage. The second example, includes unspliced, fragmentary TM additionally detected by TeloPrime.

**Figure S7.**
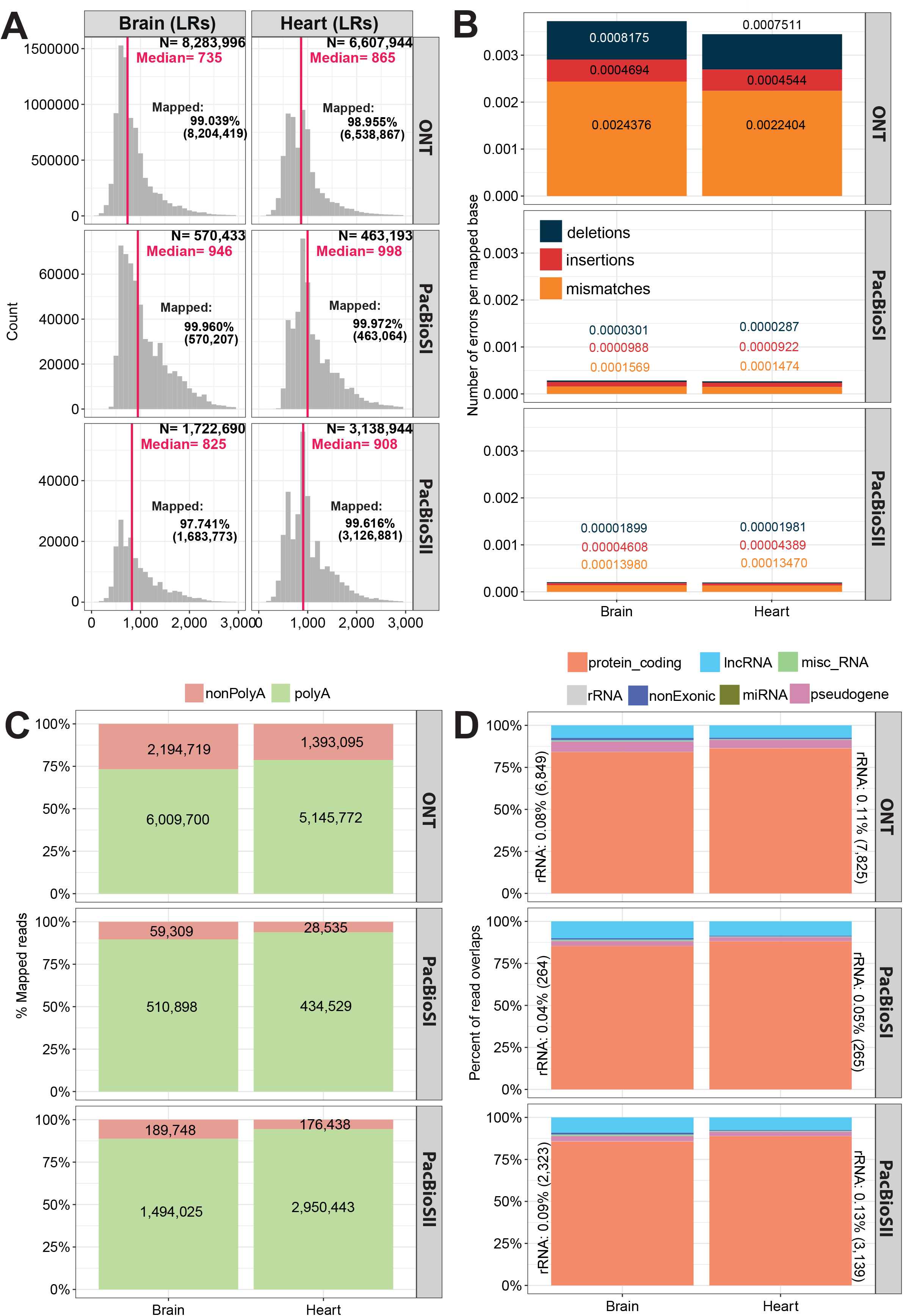
Basic long-read statistics for CapTrap-seq across different sequencing platforms. **(A)** Length distribution of raw long-read reads for ONT, PacBioSI and PacBioSII sequencing platforms. The total number of reads (N), median read length (red and vertical line) and the mapping rate are shown in the top right corner. **(B)** The number of errors per mapped bases (error rate) for CapTrap-seq for ONT, PacBioSI and PacBioSII sequencing platforms. The colour code as specified in Figure S3; **(C)** Proportion of poly(A) (green) and non-poly(A) (red) ONT and PacBio (SI and SII) reads; **(D)** The GENCODE gene biotype detection for ONT and PacBio sequencing platforms. Biotype classes as described in Figure S4. The proportion of raw reads mapping to rRNA along with absolute raw read numbers in parenthesis is given on the outer side of each bar.

**Figure S8.**
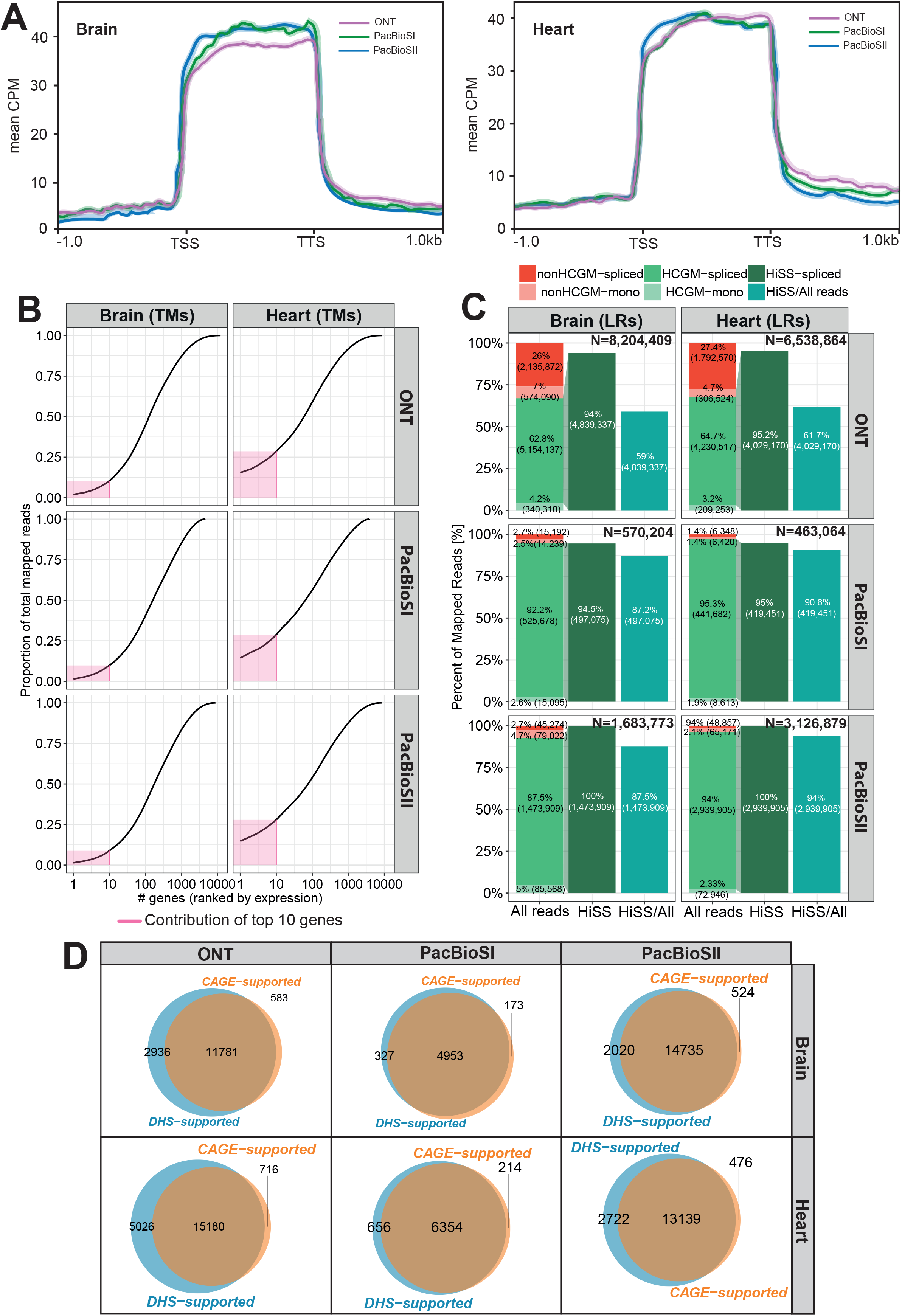
The read alignment processing and transcript model generation statistics. **(A)** Read aggregate deptools^49^ profiles along the body of annotated GENCODE genes for the CapTrap-seq combined with different long-read sequencing platforms; **(B)** Cumulative transcript curve showing the proportion of mapped reads (Y axis) captured by the top n more expressed genes (x-axis). The pink area represents the proportion of reads captured by the top ten gene expressors; **(C)** High-Confidence Genome Mapping (HCGM) and Hi-Seq-Supported read mappings (HiSS) for spliced and unspliced long reads. The stacked bar plots show the proportion of mapped reads according to HCGM classification (see legend figure S4 for more details on HCGM); **(D)** Venn diagrams showing transcripts with CAGE (orange) and DHS (blue) support for ONT and two PacBio platforms in the human heart and brain CapTrap-seq samples;

**Figure S9.**
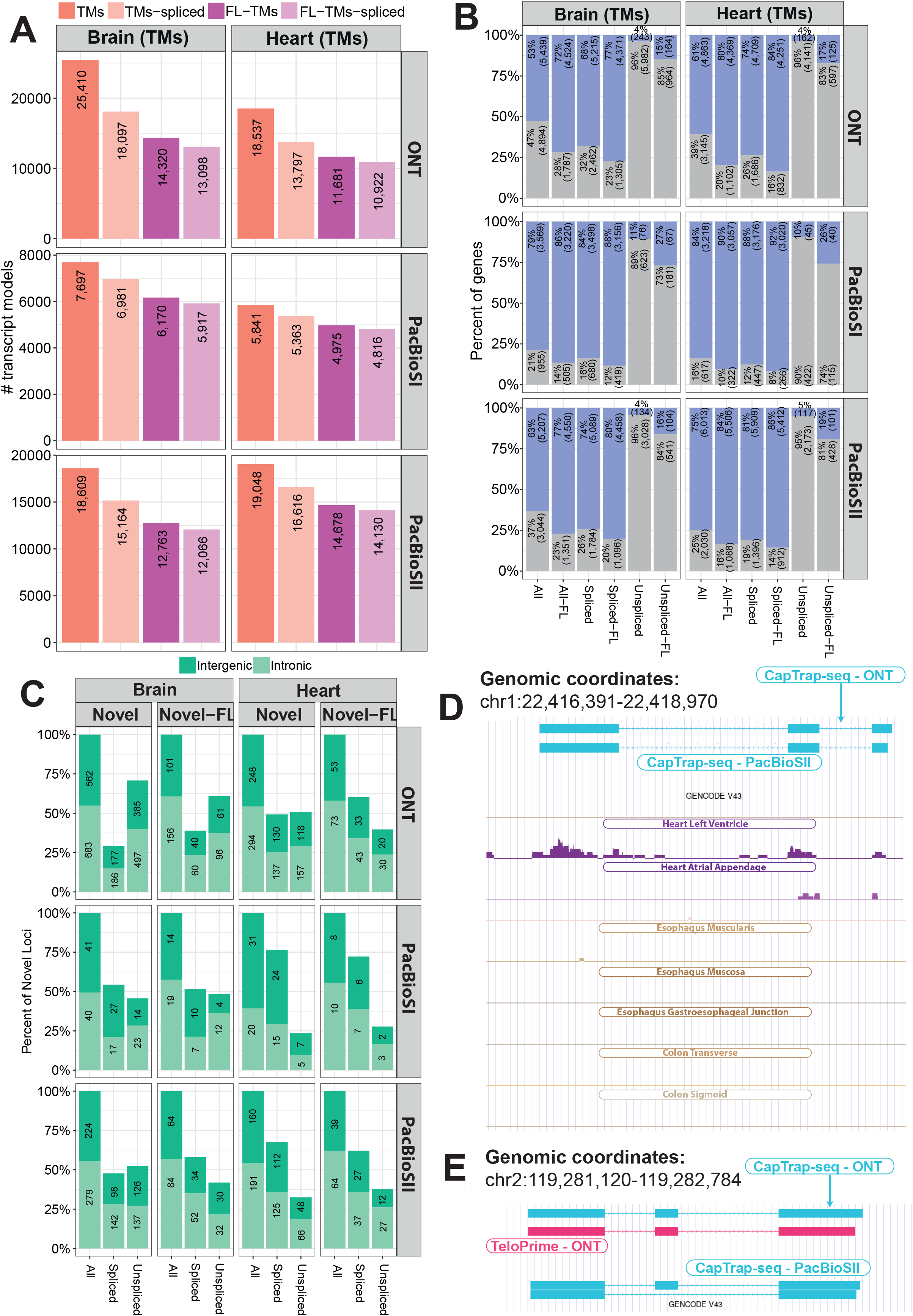
Gene detection rates for CapTrap-seq across different long-read sequencing platforms. **(A)** Bar plots showing the number of all TMs (red), spliced TMs (coral), FL-TMs (purple) and, FL-TMs-spliced (lilac); **(B)** Proportion of GENCODE (v24) genes with complete (blue) or partial (gray) detection by six different classes of identified TMs. See Figure S5B for details; **(C)** Proportion of novel and novel full-length intergenic (dark green) and intronic (light green) loci (all, spliced and unspliced in GENCODE v24) by CapTrap-seq in different long-read platforms; **(D)** A novel, heart-specific, full-length spliced locus, whose expression profile is additionally supported by GTEx^43^ RNA-seq data. **(E)** A missing alternative isoform detected by PacBioSII platform for a novel, spliced full-length brain-specific locus detected by CapTrap-seq and TeloPrime in combination with ONT sequencing (see Figure S6B for details);

**Figure S10.**
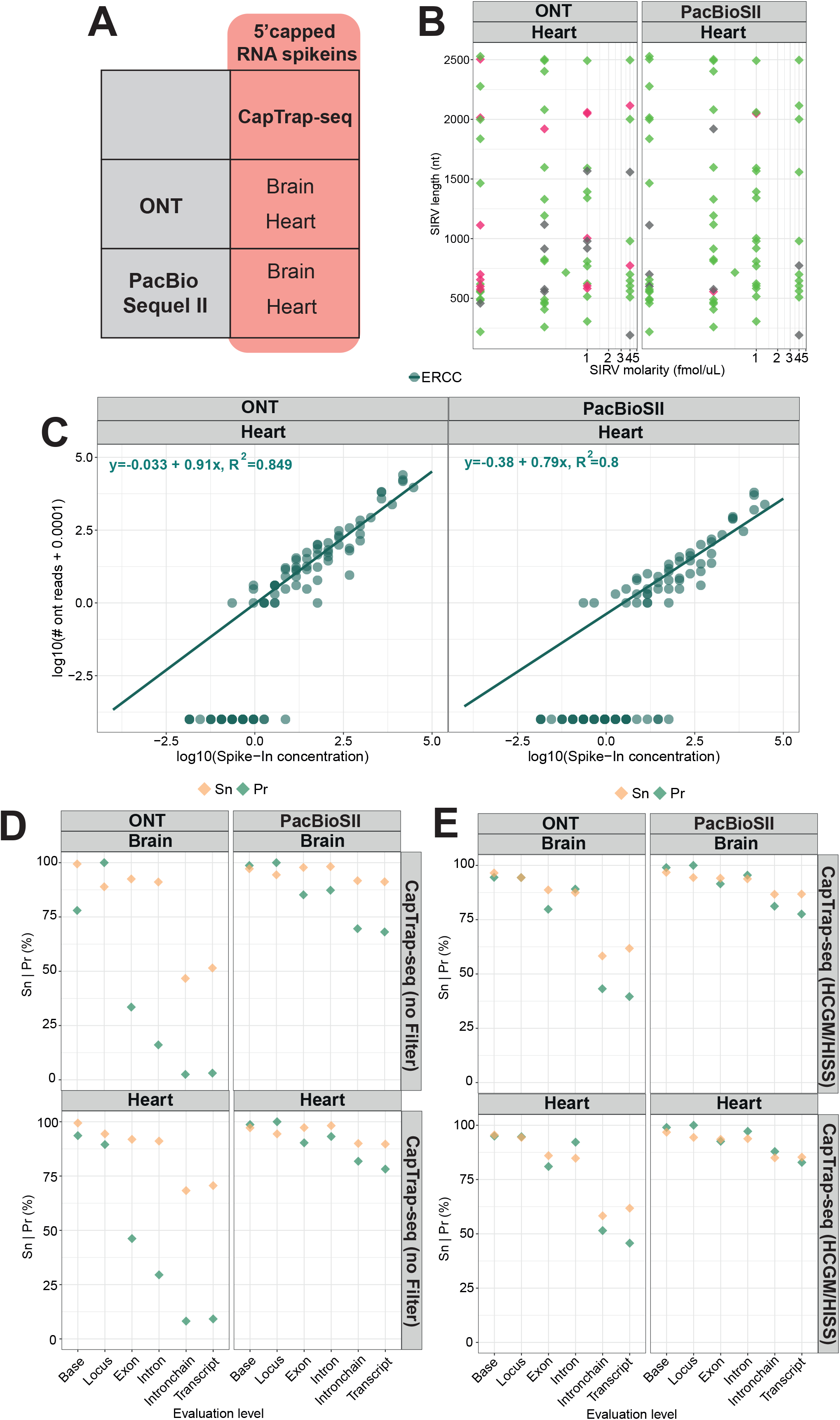
Employing synthetic RNA spike-ins to test the performance of CapTrap-seq and the fidelity of generated TMs. **(A)** Experimental design for unbiased (92 ERCC spike-ins) and targeted (42 probed and 50 non-probed ERCC spike-ins) CapTrap-seq experiments using the capped synthetic ERCC and SIRV controls; (**B**) SIRV-based transcript reconstruction accuracy for the heart sample sequenced using ONT and PacBioSII platforms. Green indicates end-to-end detection, red partial detection, while gray is the absence of a given SIRV. Detection of SIRVs is shown here as a function of initial concentration and length; **(C)** Correlation between input RNA concentration and raw read counts for ERCC spike-ins in the heart sample. The individual data points represent 92 synthetic ERCC RNA sequences (for further details see Figure 5); The analysis of sensitivity (orange) and precision (green), calculated at various levels (nucleotide, exon, intron, transcript, gene) for SIRV by gffcompare^50^ using raw **(D)** and processed (HCGM/HiSS) **(E)** alignments.

## SUPPLEMENTARY TABLES

**Table S1.**
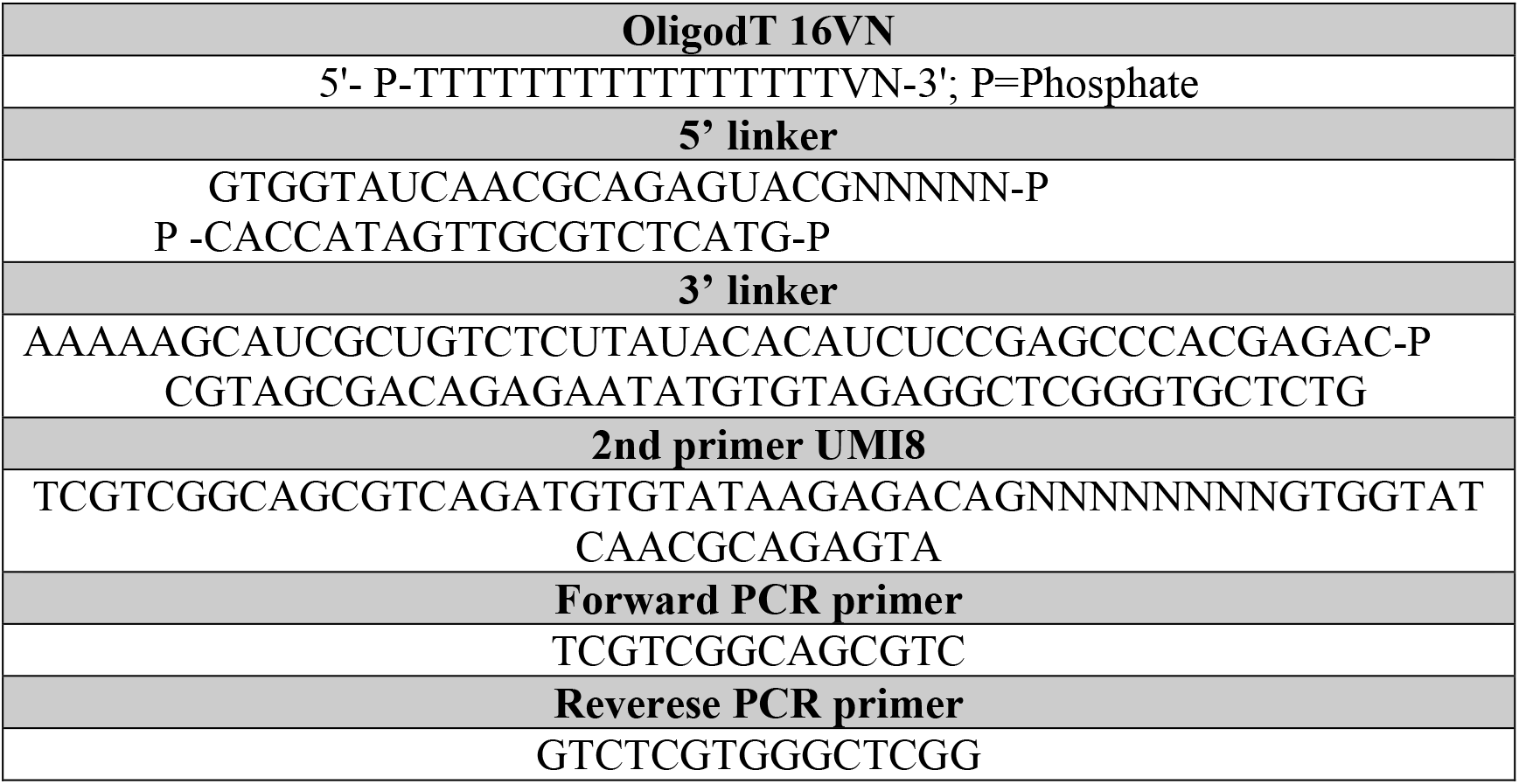
List of primers and linker sequences used in this study.

## Author contributions

B.U.-R., R.G., S.C.-S. , and J.L. designed the experiment. S.C.-S. optimized the CapTrap-seq for long-read sequencing with the input from H.N., H.T. and P.C. S.C.-S. and C.A. generated cDNA/RNA libraries and performed the direct RNA and cDNA sequencing. S.C.-S optimized the capping procedure of RNA spike-ins. J.L. developed the LyRic pipeline, *tmerge* and *buildLoci*. B.U.-R.and E.P. analyzed the data. B.U-R. and R.G. wrote the manuscript with contributions from S.C.-S. and J.L.

## Acknowledgements

We thank the Guigó laboratory for their valuable input and help with sample handling. We also thank the GENCODE experimental group for their valuable input and discussion, in particular A.Frankish, J.Mudge (EBI, UK) and Rory Johnson (UCD, Ireland), and also thanks to the entire GENCODE group, in particular, M.Diekhans (UCSC, US). We would like to acknowledge the CRG Genomics Unit for assistance with short-read Illumina sequencing and the NGS Sequencing Core facility headed by S. Goodwin for their assistance in generating PacBio Sequel I and Sequel II data.

This work and its publication were supported by the National Human Genome Research Institute of the US National Institutes of Health (grant 2U24HG007234-09) and the National Science Center PL grant (2018/31/B/NZ2/01940 to B.U.R.). We acknowledge the support of the Spanish Ministry of Science and Innovation to the EMBL partnership, Centro de Excelencia Severo Ochoa and CERCA Programme / Generalitat de Catalunya.

We thank R. Garrido and R.Carbonell Garcia (CRG), as well as A. Chmielewska and K. Solka (IBCH PAS) for administrative support.

## Data and code availability

All computer code is available from the authors upon request. The LyRic (https://github.com/guigolab/LyRic), *tmerge* (https://github.com/guigolab/tmerge) and *buildLoci* (https://github.com/julienlag/buildLoci) codes are available from the github repository. Sequence data have been deposited in the European Nucleotide Archive (ENA) repository under accession number E-MTAB-13063.

## Disclaimer

The content is solely the responsibility of the authors and does not necessarily represent the official views of the National Institutes of Health.

## Conflict of interest

The authors declare no conflict of interest.

## MATERIALS AND METHODS

### Capping ERCC and Lexogen SIRV spike-in controls

The Capping reaction was performed using Vaccinia capping enzyme (catalog num M2080S, New England BioLabs) following the recommendations of the manufacturer’s capping protocol (https://international.neb.com/protocols/0001/01/01/capping-protocol-m2080) with two changes: 3.5 µl of RNAse inhibitors (RNasin Plus RNase Inhibitor, catalog num N2611, Promega) were added to the capping reaction to avoid RNAse degradation, and the incubation time was extended from 30 minutes to two hours, following a recommendation from New England BioLabs technical support scientists. The final capping reaction was purified by using 1.8x AMPure RNA Clean XP beads (catalog num. A63987, Beckman Coulter) and resuspended in 100 μl of nuclease-free water.

### Samples and cDNA library preparation

Two total RNA human commercially available heart and brain RNA samples were used to benchmark combinations of library preparation methods and sequencing platforms.performance: total RNA from adult brain (Ambion - catalog num. AM7962; lot num. 1887911 and1997739, Thermo Fisher) and total RNA from adult heart (Ambion - catalog num. AM7966; lot num. 1866106 and 1906770, Thermo Fisher). The total RNA samples were quality controlled for concentration and integrity.

The obtained RNA was processed using four different long-read library preparation methods as indicated below:

1. **CapTrap-seq full-length cDNA libraries** The protocol starts with first strand synthesis where 5 μg of total RNA were mixed in a 10 μl reaction with 1μl of dNTPs (10 mM; catalog num. 18427013, Life Technologies) and 0.5 μl of following anchored priming oligo: OligodT 16VN (100 μM, Sigma-Aldrich) (Table S1). The mixture was incubated at 65°C for 5 minutes and immediately cooled on ice for 1 min. The enzyme mix (PrimeScript II Reverse Transcriptase, catalog num. 2690A, Takara) was prepared as follows: In a total 10 μl reaction 4 μl of 5x PSII buffer, 1 μl of RNasin Plus RNase Inhibitor (catalog num. N2611, Promega), 4 μl of nuclease-free water (UltraPure DNase/RNase-Free Distilled Water, catalog num. 10977035, Invitrogen) and 1 μl of PrimeScript II RTase (200U/μl). Ten μl of this enzyme mix were added to each RNA/primer mix reaction in a total volume of 20 μl reaction. First-strand synthesis was performed according to the following program: 42 °C for 60 minutes and held at 4°C. The resulting products were purified with 1.8x AMPure RNA Clean XP beads (catalog num. A63987, Beckman Coulter) and resuspended in 42 μl of nuclease-free water. The purified first-strand (40 μl) was then mixed with 2 μl of 1M NaOAc (pH 4.5) and 2 μl of NaIO4 (250 mM) in a total volume of 44 μl. The mixture was wrapped with aluminium foil and incubated for 5 minutes on ice to allow the oxidation of the diol group, present in the m7G cap structure, to aldehyde. Sixteen μl of Tris HCl (1M, pH 8.5) were added to the mixture to stop the oxidation reaction and the whole reaction was quickly purified with 1.8x AMPure RNA Clean XP beads and resuspended in 42 μl of nuclease-free water. The aldehyde groups resulting in a 40 μl mix were then biotinylated using a mixture containing 4 μl of NaOAc (1M, pH 6.0) and 4 μl of Biotin (Long Arm) Hydrazide (100 mM, catalog num. SP-1100, Vector Laboratories). The 48 μl resulting mixture was next incubated for 30 minutes at 40°C, purified with 1.8x AMPure RNA Clean XP beads, and resuspended in 42 μl of nuclease-free water. The RNase ONE Ribonuclease (catalog num. M4261, Promega) was used at this point to eliminate the Biotin groups at the 3’ end by degrading only the single-strand RNA present into the mixture. Only the biotin bound to m7G cap structure at 5’end was kept during this step, where 0.5 μl of RNase ONE enzyme (10 U/μl) and 4.5 μl of 10x RNase ONE Buffer were added to the 40 μl of biotinylated product. The resulting mixture was incubated for 30 minutes at 37°C and afterwards, purified with 1.8x AMPure RNA Clean XP beads and resuspended in 42 μl of nuclease-free water. The m7G cap structure bound to biotin is then selected using M-270 streptavidin magnetic beads. Before starting, 30 μl of M-270 streptavidin magnetic beads (catalog num. 65305, Thermo Fisher Scientific) were washed twice with 30 μl of CapTrap LiCl binding buffer (c3.64 ml of nuclease-free water, 35 ml of Lithium chloride 8M, 0.8 ml of 1M Tris-HCl at pH 7.5, 0.4 ml of 10% Tween 20 and 0.16 ml of 0.5M EDTA at pH 8.0) and resuspended with 95 μl of same LiCl binding buffer. The 40 μl of the sample recovered after RNase ONE purification were mixed with 95 μl of washed M-270 streptavidin magnetic beads and incubated at 37°C for 15 minutes. During the incubation time, the reaction was mixed by pipetting 10 times at a 7-minute interval to ensure that the beads remain in suspension. Beads were pulled down with a magnet and after supernatant removal, they were washed 3 times with 150 μl of CapTrap washing buffer (39.12 ml of nuclease-free water, 0.4 ml of 1M Tris-HCl at pH 7.5, 0.4 ml of 10% Tween 20 and 0.08 ml of 0.5 M EDTA at pH 8.0). After washing steps, the single strand cDNA was released using 35 μl of CapTrap release buffer (1x RNase ONE buffer with 0.01% Tween 20), heat shocked for 5 minutes at 95°C, and quickly cooled on ice. The supernatant was then recovered from M-270 streptavidin magnetic beads and stored on ice; meanwhile, a second release was performed adding 30 μl of CapTrap release buffer and the supernatant was also collected and mixed with the first release eluate. The final 65 μl volume was treated with 5 μl of an enzymatic mixture containing: 0.1 μl of RNase H (60 U/μl, Ribonuclease H <RNASE H>, catalog num. 2150, Takara), 2 μl of RNase ONE (10 U/μl) and 2.9 μl CapTrap release buffer, incubated for 30 minutes at 37 °C and afterwards, purified with 1.8x AMPure XP beads (catalog num. A63881, Beckman Coulter) and resuspended in 42 μl of nuclease-free water. The purified sample (approx 40 μl) was concentrated using a speed vac, by drying it for 35 min at 80 °C. The dried sample is resuspended with 4 μl of nuclease-free water. Two different linkers (Sigma-Aldrich), a 5’ linker and a 3’ linker, were ligated to the single-stranded cDNA using a 2-step linker ligation^51^ (Table S1). During the first linker ligation, 1 μl of 5’ linker (10 μM) was ligated with 10 μl of Mighty mix (DNA Ligation Kit <MIGHTY Mix>, catalog num. 6023, Takara) to 4 μl of the sample coming from the previous step. Just before proceeding with ligation incubation (4 hours at 30°C or 16 hours at 16°C), the linker and the sample are denatured separately for 5 min at 55°C and 95 °C respectively, put on ice for two minutes and then added to the Mighty ligation mix. The 5’ linker ligation product was purified twice, to completely eliminate the non-incorporated linkers, with 1.8x AMPure XP beads and finally resuspended in 42 μl of nuclease-free water. The resuspended sample (approx 40 μl) was concentrated using a speed vac, by drying it for 35 min at 80 °C. The dried sample was resuspended with 4 μl of nuclease-free water. During the second linker ligation, 1 μl of 3’ linker (10 μM) was ligated with 10 μl of Mighty mix to 4 μl of the sample coming from the 5’ linker ligation step. Just before proceeding with ligation incubation (4 hours at 30°C or 16 hours at 16°C), the linker and the sample were denatured separately for 5 min at 65°C and 95 °C respectively, put on ice for two minutes and then added to the Mighty ligation mix. The 3’ linker ligation product was purified with 1.8x AMPure XP beads and finally resuspended in 42 μl of nuclease-free water. The double-stranded linkers were converted into single-stranded to successively allow the second strand synthesis by Shrimp Alkaline Phosphatase (1 U/μl SAP, catalog num. 78390, Affymetrix) and Uracil-Specific Excision Reagent (1 U/μl USER, catalog num. M5505L, NEB) treatment. The 40 μl sample final volume was combined with 2 μl of nuclease-free water, 5 μl of 10x SAP buffer/10xTE, 1 μl of SAP (1 U/μl), and 2 μl of USER (1 U/μl), mixed, incubated for 30 minutes at 37°C, 5 minutes at 95°C and finally placed on ice. Afterwards, the sample was purified with 1.8x AMPure XP beads and finally resuspended in 42 μl of nuclease-free water. The samples were concentrated using a speed vac (40 minutes at 80°C) and afterwards resuspended with 5 μl of nuclease-free water, before second strand synthesis. 20 μl of second strand synthesis mix, prepared with 5.8 μl of nuclease-free water, 12.5 μl of 2x HiFi KAPA mix (catalog num. 7958927001-KK2601, Kapa), 0.5 μl of 2nd primer UMI8 (Table S2, 100μM, Sigma-Aldrich), containing 8 mer UMI sequence and 1.3 μl of DMSO, were added to 5 μl of the sample. The mixture was incubated for 5 minutes at 95°C, 5 minutes at 55°C, 30 minutes at 72°C and finally held at 4°C until 1 μl Exonuclease I (20U/μl, catalog num. M0293S, NEB) was added to each sample. The sample was then incubated for 30 minutes at 37°C and afterwards, purified twice with 1.8x and 1.4x, respectively AMPure XP beads and finally resuspended in 42 μl of nuclease-free water. The samples were dried up with speed-vac (75 minutes at 37°C) and resuspended with 5 μl of nuclease-free water. These 5 μl were used to amplify the cDNA CapTrap library by Long and accurate PCR (LA-PCR). In the way to avoid PCR replicates these five μl of each sample were split into two PCR-independent reactions. The PCR reaction mix (TaKaRa LA Taq, catalog num. RR002M, Takara) was assembled using 24 μl of nuclease-free water, 5 μl of 10x Buffer, 5 μl of MgCl2 (25 mM), 8 μl of dNTPs mix (2.5 mM each), 2.5 μl of each primer (Table S1), 0.5 μl of La TaKaRa Taq (5 U/μl) and 2.5 μl of second strand synthesis product, with a final volume of 50 μl. The PCR cycling conditions used were 30 seconds at 95°C for the denaturation step, 16 cycles of 15 seconds at 95°C, 15 seconds at 55°C and 8 minutes at 65°C for amplification steps, followed by 10 minutes at 65°C and hold at 4°C. The 2 PCR replicates were merged and purified with 1x AMPure XP beads and finally resuspended in 22 μl of nuclease-free water. Samples were quantified with Qubit (Qubit 4 Fluorometer, Thermo Fisher Scientific) and quality checked with BioAnalyzer (Agilent 2100 Bioanalyzer, Agilent Technologies).
2. **SMARTer Full-length cDNA libraries** We used an aliquot of 4 ug total RNA for each sample. Each aliquot of 4 μg was depleted from ribosomal RNA with 10 μl of rRNA removal solution from the Ribo-Zero kit (RiboZero kit, catalog num. MRZH11124, Epicentre-Illlumina), strictly following the manufacturer’s protocol. A total of 3.5 μl ribo-depleted material was used for SMARTer (SMARTer PCR cDNA Synthesis Kit, catalog num. 634926 and Advantage 2 PCR kit, catalog num. 639207, Clontech-Takara) protocol. Libraries were prepared strictly following the manufacturer’s protocol.
3. **Teloprime Full-length cDNA libraries** 2 μg of each total RNA sample were used to start the Teloprime cDNA library preparation strictly following the manufacturer’s protocol.
4. **Direct RNA ONT libraries** The total RNA samples were first poly(A) enriched using Dynabeads® Oligo (dT) following manufacturer protocol. Two rounds of poly(A) enrichment were performed to clean and recover maximum amounts of poly(A) RNA. The full protocol was performed using LiDS as a detergent to prepare washing, lysis, and binding buffers. We repeated the poly(A) enrichment procedure several times, depending on the quality of the RNA sample, until we obtained a total amount of 500 ng of poly(A) RNA. Next, we proceeded with library preparation using the SQK-RNA002 kit. The T4 DNA ligase was used with Quick Ligation Reaction Buffer to anneal and ligate the Direct RNA RT adapters (RTA oligo) to the sample. To create a cDNA-RNA hybrid, adapter-ligated poly(A) RNA was incubated with dNTPs, 5x first-strand buffer, nuclease-free water, and Maxima RT enzyme was used instead of SuperScriptIII to ensemble the reverse transcription reaction. Tubes were placed in a thermal cycler and incubated for 1h at 60°C and 5 minutes at 85°C. The reverse-transcription step is highly recommended by the Nanopore protocol to obtain high sequencing throughput of direct RNA samples because it reduces RNA secondary structures and gives stability to the RNA molecule before it passes through the pore and is sequenced. RNAClean XP beads were used to purify the RT reaction and finally, the sequencing adapters are ligated to the RNA/cDNA hybrid by using NEBNext Quick Ligation Reaction Buffer and T4 DNA Ligase following the manufacturer’s protocol. In the last purification step, RNAClean XP beads were employed to clean RNA after the adapter-ligation reaction to make it ready for Qubit quantification and fragment distribution quality check.

### ONT and PacBio sequencing

The CapTrap cDNA libraries were split into two different aliquots to meet the sequencing platform requirements. Next, platform-specific sequencing libraries were prepared and loaded into ONT and PacBio long-read sequencing platforms. The TeloPrime, SMARTer and direct RNA libraries were only sequenced using the ONT platform.

The ONT cDNA sequencing was performed with 500-1000 ng of cDNA sample coming from cDNA-based protocols (CapTrap, SMARTer, and Teloprime) and strictly following the ONT SQK-LSK108 and SQK-LSK109 adapter ligation protocols. Direct RNA sequencing was performed with 500 ng of poly(A) RNA. For the ONT sequencing, we used a MinION device, ONT R9.4 flow cells and the standard MiniKNOW protocol script.

PacBio Sequel and Sequel II cDNA sequencing was performed using 500 ng of cDNA samples and following the general protocol workflow for amplicon sample preparation and sequencing SMRTbellTM Express Template Prep Kit 2.0. This workflow allows the generation of highly accurate sequences from amplicons ranging in size from several hundred bases to 10 kb or larger. The samples were prepared following the procedures and details specific to the preparation of cDNA libraries recommended by the manufacturer.

None of the RNA samples and cDNA libraries were subjected to size selection before loading them on ONT and PacBio Sequel (I and II) platforms.

### Illumina Short read SMARTer sequencing

We produced matched short-read RNAseq data in the human and brain samples to be used by LyRic to support long reads and to generate the HiSS. A total of 250 ng of each cDNA (previously prepared for long-read cDNA libraries) were used as starting material. The samples were fragmented using Covaris under the following parameters: 10% Duty cycle, Intensity 5, 200 Cycles per burst, during 30 seconds in a total volume of 55 μl. Illumina sequencing library was prepared using a TruSeq-based protocol (KAPA LTP Library Preparation kit Illumina, catalog num. 7961898001, Roche) and manufacturer protocol was strictly followed. All samples were barcoded and multiplexed using the Illumina barcoding system (6 mers) and then sequenced in a Hi-Seq 2500 lane with HiSeq Sequencing v4 Chemistry. A mean of 22.9 million 125-base paired-end reads was generated for each sample. Paired-end reads were then checked for quality and processed together with long-reads by the LyRic pipeline.

### Data analysis with the LyRic pipeline

LyRic is a versatile automated transcriptome annotation and analysis workflow written in the Snakemake language^32^. The RNA sequencing ONT, PacBio and Illumina reads were mapped to the human reference genome assembly GRCh38/hg38 (in addition to sequences of 96 ERCC and 69 SIRV Spike-In Controls). We used Minimap2^33^ and STAR (v2.7.6a)^48^ to map long and short reads to the genome, respectively and a custom reference gene annotation file was used to compare the LyRic output and the annotation. A custom reference gene annotation file was built by combining GENCODE gene annotation (v24), SIRV annotation containing 69 transcripts and 92 ERCC spike-in controls. The read aggregate profiles along the body of annotated GENCODE genes were generated using the deptools2^49^ package. The computeMatrix and plotProfile functions are set as described in the LyRic documentation.

The read-to-genome alignments were used to identify High Confidence Genome Mappings (HCGMs). The HCGMs are the read-to-genome mappings characterized by four main features, including (1) the presence of only canonical introns (spliced reads only), (2) no suspicious introns possibly arising from RT template switching (spliced reads only), (3) minimum average sequencing quality around the splice junctions (spliced reads only) and (4) presence of polyA tail in the read sequence (for unspliced reads only). Next, we identified Hi-Seq-Supported read mappings (HiSS), which correspond to HCGMs that have all their splice junctions supported by at least one split read in the corresponding short-read Illumina HiSeq sample. The short-read Illumina HiSeq data generated using the SMARTer protocol was used to support the ONT reads, while the PacBio ones were processed without short-read support. Thus, the HiSS PacBio reads are exactly equivalent to HCGMs. Finally, HiSS were merged into a non-redundant set of transcript models (TMs) using *tmerge*.

Next, we merged the LyRic TMs with the GENCODE annotation (v24) using *tmerge*, identified novel loci using *buildLoci* and classified them as intronic and intergenic. We selected the GENCODE v24 set to make the output of this analysis compatible with the findings of the Long-read RNA-seq Genome Annotation Assessment Project (LRGASP) Consortium^24^.

Full LyRic documentation is available at https://guigolab.github.io/LyRic/documentation.html, while the LyRic data portal with detailed information, including the config file is available at https://github.com/guigolab/captrap-seq.

### Estimating CapTrap-seq sensitivity, quantitativeness and TM accuracy using spike-ins

The ERCC spike-in mix 1 and Lexogen SIRV_Set 1 mix (generated using 1:1 E1 and E2 mixes to get the maximum dynamic range) using 1:1 proportion ERCCs:SIRVs were added to each sample before the per library preparation. The analysis of the correlation between the initial ERCC spike-in concentrations and raw long-read counts (Figure 5D and S10C) shows a detection limit ∼8.4x10-2 attomol (-1.875 in log10 units) for sequenced molecules. This threshold is equivalent to 2,016 molecules, as in 4 μg of a 1:100 dilution of ERCC spike-in was added to 4 μg of each RNA sample. This value approximately equals 1.05x10-2 molecules per cell on the assumption that the total RNA content of a single cell is 5 pg^52^.

To evaluate the accuracy of SIRV transcript reconstruction, we used a complete SIRV annotation containing all 69 SIRV transcripts. The obtained SIRV transcript models were compared against the reference SIRV annotation using gffcompare^50^. The SIRV-Set 1 annotations are available at https://www.lexogen.com/sirvs/download/.

## Notes

### Competing Interest Statement

The authors have declared no competing interest.

